# A novel RAB11-containing adaptor complex anchoring myosin-5 to secretory vesicles

**DOI:** 10.1101/2021.09.12.459962

**Authors:** Mario Pinar, Ana Alonso, Vivian de los Ríos, Ignacio Bravo-Plaza, Álvaro Gandara, Ernesto Arias-Palomo, Miguel Á. Peñalva

## Abstract

Hyphal fungi grow rapidly by apical extension, providing a notorious example of polarized growth. The continuous supply of secretory vesicles necessary to meet the demands of the extending tip and the long intracellular distances existing between the tip and the basal septum, often localized > 100 µm away from the former, impose the need of efficient networks of intracellular traffic involving exquisite cooperation between microtubule- and actin-mediated transport. In *Aspergillus nidulans* kinesin-1 conveys secretory vesicles to the hyphal tip, where they are transferred to myosin-5, which focuses them at the growing apex, thereby determining cell shape. This relay mechanism and the central role played by myosin-5 in hyphal morphogenesis suggested that the mechanisms anchoring secretory vesicles to this motor should involve specific adaptor(s) ensuring the robustness of actomyosin-dependent transport.

Secretory vesicles are charged with RAB11, a regulatory GTPase that determines the Golgi to post-Golgi identity transition. By using a combination of shotgun proteomics, GST-RAB pull-down assays, *in vitro* reconstitution experiments, targeted reverse genetics and multidimensional fluorescence microscopy with endogenously tagged proteins we show that RAB11, the master regulator of fungal exocytosis, mediates myosin-5 engagement both by contacting the motor and by recruiting UDS1, a homologue of an as yet uncharacterized *Schizosaccharomyces* protein ‘upregulated during mitosis’, which we demonstrate to be a novel RAB11 effector. Analytical ultracentrifugation determined that UDS1 is an elongated dimer and negative-stain electron microscopy showed that, in agreement, UDS1 is rod-shaped. UDS1 does not contact myosin-5 directly, but rather recruits the coiled-coil HMSV, which bridges RAB11/UDS1 to myosin-5. An HMSV-scaffolded complex containing UDS1 and myosin-5 is present in cells, and a RAB11-UDS1-HMSV complex can be reconstituted *in vitro* in a RAB nucleotide state-dependent manner. In the absence of UDS1/HMSV the steady state levels of myosin-5 at the apical vesicle supply center diminish markedly, such that microtubule-dependent transport spreading vesicles across the apical dome predominates over apex-focused actin-mediated transport. As a consequence, RAB11 and chitin-synthase B (a cargo of the RAB11 pathway) are not focused at the apex, being distributed instead across the apical dome. Therefore, the RAB11 effector UDS1/HMSV cooperates with the GTPase to adapt secretory vesicles to myosin-5, which is required for the apical targeting of RAB11 cargoes and thus for the normal morphology of the hyphae.

## Introduction

How the multiplicity of membranous cargoes of eukaryotic cells are specifically adapted to molecular motors constitutes a fundamental question of cell biology. Across the eukaryotic realm, type V myosins play a key role in the transport of these cargoes, often acting in concert with microtubule-dependent motors (Hammer & Sellers, 2012). For example, in the filamentous fungus *Aspergillus nidulans*, a single myosin-5 (denoted MyoE) and a kinesin-1 (KinA) cooperate to transport RAB11 secretory vesicles (SVs) originating at the Golgi to the hyphal apices (Pantazopoulou et al., 2014, Peñalva et al., 2017).

In *A. nidulans* and other hyphal fungi these SVs concentrate at an apical structure denoted Spitzenkörper (SPK), which acts as a vesicle supply center from which SVs are delivered to the growing tip’s plasma membrane. The SPK contains a F-actin organizing center (Sharpless & Harris, 2002), such that actin cables span the region of the tip spreading out from the apex like the ribs of an umbrella. In contrast, microtubules (MTs) make apical contacts with their plus-ends at a broader, crescent-shaped region of the tip denoted ‘the apical dome’.

A division of roles underlies cooperation between actomyosin and microtubule (MT) transport in *A. nidulans* (Pantazopoulou et al., 2014, Peñalva et al., 2017, Pinar et al., 2015, Pinar & Peñalva, 2020, Schuchardt et al., 2005, Zhang et al., 2011): kinesin-1 conveys RAB11 SVs to the hyphal tips whereas myosin-5 concentrates them at the SPK (Figure 1A). The partially redundant role played by kinesin-1 makes myosin-5 non-essential, although its absence slows down growth markedly and causes morphological abnormalities resulting from inability to focus exocytosis at the apex. Cooperation between the microtubule and the actin cytoskeletons is not uncommon in tip-growing cells of organisms that are evolutionary distant from fungi. Another notable example of this cooperation occurs in the protonema of the moss *Physcomitrella patens*, which contains a cluster of F-actin at the apex that governs the directionality of growth, and that strikingly resembles the fungal SPK/vesicle supply center (Wu & Bezanilla, 2018)

**Figure 1.**
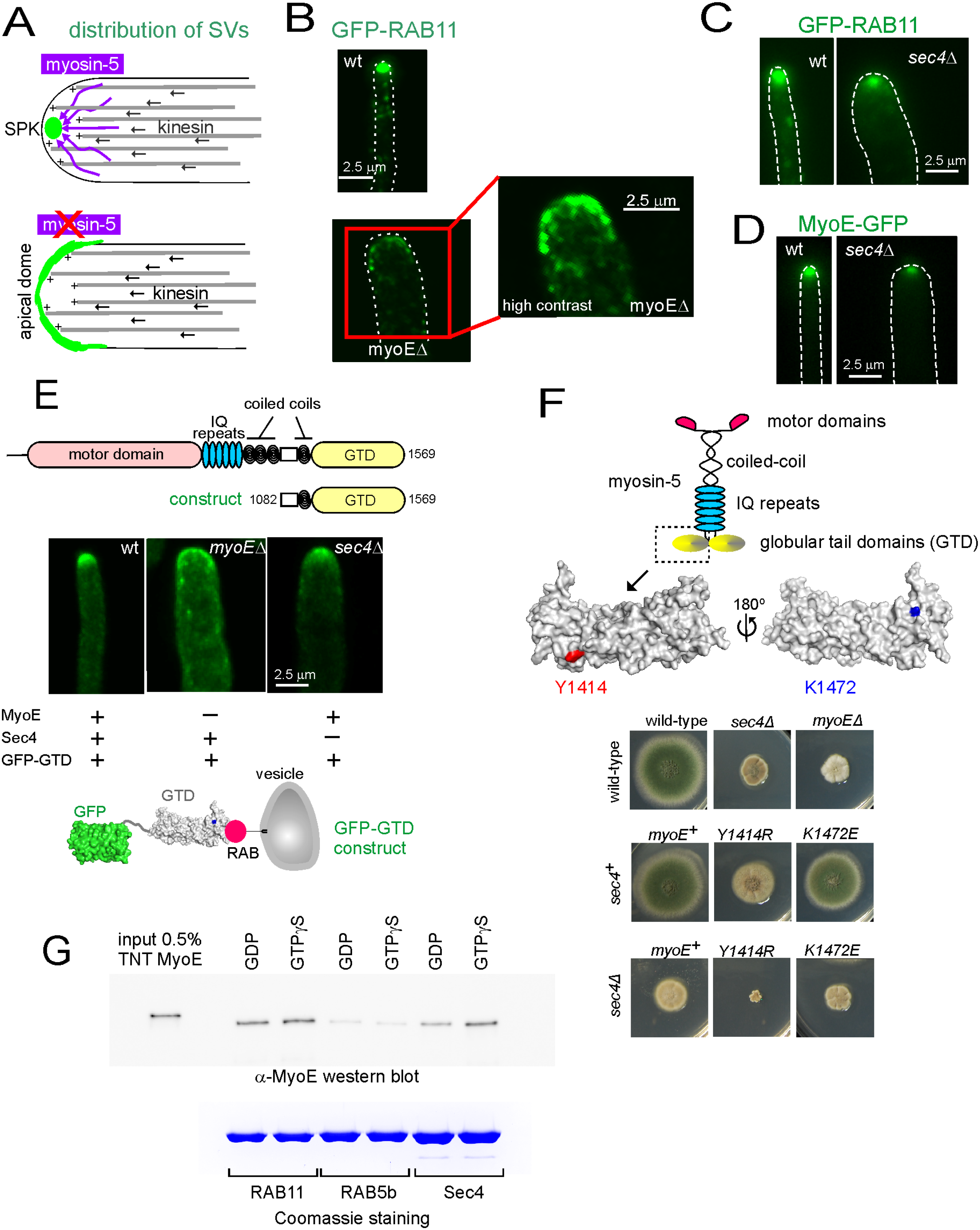
SVs can be delivered to the vesicle supply center by myosin-5 in the absence of Sec4: myosin-5 is a RAB11 effector. (A) Schemes depict the cooperation of kinesin-1 and myosin-5 to deliver RAB11 SVs to the SPK and the situation in a myosin-5-less mutant in which SVs distribute across the apical dome because they cannot be focused at the SPK due to the absence of F-actin-dependent transport. (B) Localization of GFP-RAB11 in the wild-type and in a *myoE*Δ mutant lacking myosin-5, displayed with the same contrast adjustments. On the right image the magnified *myoE*Δ tip on the left has been contrasted to reveal better the localization of RAB11 SVs in this mutant. (C) Localization of GFP-RAB11 in the wild-type and in a *sec4*Δ tip shown at equivalent contrast. (D) Localization of MyoE-GFP in the wild-type and in a *sec4*Δ tip shown at equivalent contrast. (E) Localization of GFP-GTD expressed under the control of the sucrose-inducible *inuA* promoter (Hernández-González et al., 2018b) from a single copy transgene integrated at the *inuA* locus. The construct lacks the motor domains, the IQ repeats and most of the coiled-coil regions of MyoE (scheme on the top). Wild-type and *sec4*Δ cells contain intact the resident copy of *myoE*, whereas the *myoE*Δ cell does not, as indicated with crosses at the scheme at the bottom. (F) Localization of residues Y1414 and K1472 on opposite surfaces of the GTD (left) and growth tests (bottom) revealing that *myoE^Y1414R^*, but not *myo^R1472E^*, shows a nearly lethal negative synthetic interaction with *sec4*Δ. *myoE* missense mutant strains were constructed by gene replacement. (G) GST pull-down experiments with indicated RABs as baits, loaded with GDP or GTP*γ*S. The prey was MyoE (myosin-5) expressed in a TNT*®* coupled transcription-translation reaction. Material pulled-down by the beads was analyzed by western blotting using anti-MyoE antiserum. The bottom row shows a Coomassie-stained gel of GST-RABs used in these pull-down assays.

Membranous cargoes attach to the globular C-terminal domain (GTD) of myosin-5 *via* ‘receptors’ that are cargo-specific adaptors (Hammer & Sellers, 2012, Pashkova et al., 2006, Wu et al., 2002). In the case of SVs these adaptors contain a RAB GTPase, be it RAB11, Sec4/RAB8 or both (Wong & Weisman, 2021). In *S. cerevisiae*, Ypt31/32 (yeast RAB11s) and Sec4 (yeast Rab8) bind directly and without involvement of any other proteinaceous co-adaptor to the GTD of the myosin-5 Myo2p (Jin et al., 2011, Lipatova et al., 2008, Santiago-Tirado et al., 2011), although the levels of PtdIns4P on SVs are also important for the Myo2p-SV association (Santiago-Tirado et al., 2011). However, this model of the RAB as the only component of the myosin-5 adaptor to RAB11 vesicles is far from being universal. For example, in mammalian cells the RAB11a effector RAB11-FIP2 (RAB11 family interacting protein 2) acts as co-adaptor cooperating with the GTPase to recruit myosin-Vb to recycling endosome vesicles (Hales et al., 2002, Schafer et al., 2014, Wang et al., 2008), and in flies a protein trio consisting of myosin V, RAB11 and dRip11 deliver exocytic vesicles to the rhabdomere base (Li et al., 2007).

Intracellular distances in hyphal tip cells are remarkably large (up to 125 µm from tip to septum). Thus, it is unsurprising that *A. nidulans* uses MTs for the long-distance shuttling of membranous organelles. This feature has been experimentally advantageous to study adaptors by which organelles engage motors. For example, studies on the MT-dependent movement of early endosomes in *A. nidulans* led to the discovery of the FTS/Hook/FHIP (FHF) complex serving as adaptor between dynein and endosome cargo (Bielska et al., 2014, Qiu et al., 2019, Yao et al., 2014, Zhang et al., 2014)

Hyphae of *A. nidulans* grow by apical extension at *∼*1 µm/min at 28°C, implying that transport of SVs to the extending tip is optimized to meet the high demand of lipids that fuel the expansion in membrane surface, as well as to deliver enzymes that modify the cell wall to facilitate growth. SVs are loaded with three motors: myosin-5, kinesin-1 and dynein (Peñalva et al., 2017). It has been suggested that SVs are handed over from by kinesin-1 to myosin-5 in the region of the tip, hypothetically by switching from MT to actin cables, yet the mechanism by which myosin-5 prevails over kinesin-1 in the tip region is not understood. In view of the crucial role that myosin-5 plays in their lifestyle, we hypothesized that hyphal fungi have an adaptor by which SVs engage this motor very robustly, to ensure the efficiency of the latest step in their transport. Here we report the molecular composition of a novel adaptor that engages SVs with myosin-5. We show that myosin-5 is recruited to SVs via a RAB11 protein complex also containing UDS1 and HMSV, two proteins whose homologues in *Neurospora crassa* have been recently identified as components of the SPK (Zheng et al., 2020). Trafficking of RAB11 SVs to the SPK/vesicle supply center is impaired if this complex is disrupted, as expected for a bona fide co-adaptor of myosin-5.

## Results

### Myosin-5 is key for delivering RAB11 secretory vesicles to the hyphal tips

The efficiency of myosin-5 transport is reflected in the distribution of RAB11 SVs accumulating in the tips before fusing with the PM. In the wild-type, these SVs gather at the SPK/vesicle supply center. In *myoE*Δ cells completely lacking myosin-5 transport SVs cannot be focused at the SPK, yet they still arrive at the tip by kinesin-1/microtubule-mediated transport (Pantazopoulou et al., 2014, Peñalva et al., 2017) (Figure 1B). Consequently, RAB11 is delocalized from the SPK to a tip crescent that reflects the steady-state distribution of the microtubules’ plus-ends at the apical dome cortex (Figure 1A). This delocalization is paralleled by a conspicuous reduction of RAB11 in the tip (Figure 1B), strongly suggesting that myosin-5 is a major contributor to the transport of RAB11. Consistent with a secretory defect, loss of myosin-5 results in abnormal hyphal morphogenesis [Figure 1B, note that exocytosis determines the shape of the cell wall and markedly reduces growth (Figure 1F)(Peñalva et al., 2017, Taheri-Talesh et al., 2012)].

A candidate to adapt myosin-5 to RAB11 SVs is the RAB GTPase Sec4 acting downstream of RAB11 during transport between the TGN and the PM (Jin et al., 2011). This could be tested directly because Sec4 is not essential in *A. nidulans*, despite of its absence being nearly as debilitating as that of MyoE/myosin-5 (Figure 1F). Indeed, the amounts of RAB11 SVs accumulating in the tip were noticeably decreased in *sec4*Δ hyphae (Figure 1C). However, contrasting with *myoE*Δ mutants, *sec4*Δ mutants were still able to gather RAB11 SVs at the SPK. In agreement, myosin-5/MyoE still concentrated in the SPK in the absence of Sec4, albeit less efficiently as well (Figure 1D and Movie 1). Therefore, these data establish that there must be another adaptor sharing with Sec4 the ability to engage SVs to myosin-5. Previous studies with fungal and metazoan cells pointed to RAB11 as the most likely candidate (Goldenring, 2015, Hales et al., 2002, Lipatova et al., 2008, Roland et al., 2011).

### Both RAB11 and Sec4 interact directly with myosin-5

In the intensively studied transport of SVs to the growing bud of *S. cerevisiae*, the RAB11 homologues Ypt31/32 and Sec4 recruit Myo2 through direct binding to an amino acid patch located in the highly conserved globular tail domain (GTD) of this myosin-5 (Jin et al., 2011, Lipatova et al., 2008). If this mechanism were conserved in *Aspergillus*, the MyoE GTD domain, isolated from the rest of the protein, should bind the RABs present on the SVs, being transported with them to the tips. To test this prediction, we expressed a construct consisting of the GFP-tagged MyoE GTD domain in *Aspergillus* wild-type, *myoE*Δ and *sec4*Δ hyphae (Figure 1E). In the wt, GFP-GTD, although partly cytosolic, was present in SVs accumulating at the tip, indicating that the GTD is indeed sufficient to localize to SVs *in vivo*. This recruitment of the GTD to SVs did not depend on interaction with resident myosin-5 because in *myoE*Δ cells GFP-GTD localized to the apical dome (Figure 1C), recapitulating the distribution of RAB11 SVs (Figure 1B,C)(Movie 2). We concluded that the MyoE GTD is sufficient to bind to SVs. Remarkably, the MyoE GTD also concentrated at the apex of *sec4*Δ cells (Figure 1C), further confirming that myosin-5 transport of SVs is still operative without Sec4.

In the budding yeast, critical residue Tyr1415 is at the center of a Myo2 GTD patch that binds the RABs linking the motor to SVs (Figure 1F, schematics). Because the budding yeast does not use microtubules to transport SVs, Y1415R substitution affecting a residue crucial for the interaction between the RABs and the myosin-5 is lethal (Lipatova et al., 2008). K1473 located on the opposite GTD surface to Y1415 (Figure 1F) belongs to a patch of residues that has been reported to bind the Sec15 subunit of the exocyst (Jin et al., 2011) and to participate in GTD-motor domain interactions maintaining Myo2p in a closed conformation (Donovan & Bretscher, 2015). To investigate RAB/MyoE interactions we introduced substitutions Y1414R and K1472E (equivalent to *S. cerevisiae* Y1415 and K1473) into MyoE by replacing the wt *myoE* locus with the corresponding mutant alleles. Y1414R resulted in a conspicuous growth defect, confirming that MyoE Y1414 plays an important physiological role. In contrast K1472E caused a minor colony growth defect by itself. In double mutants *myoE (K1472E)* did not weaken *sec4*Δ strains any further. In sharp contrast, the double *sec4*Δ *myoE (Y1414R)* mutant combination was nearly lethal (Figure 1F). As Y1414 is crucial for RAB binding, these data provide strong genetic evidence that an exocytic RAB other than Sec4 is capable of binding directly to the myosin-5 GTD.

Therefore, we investigated the possibility that MyoE is a direct effector of both Sec4 and RAB11. As MyoE is insoluble when expressed in bacteria, we synthesized MyoE in a coupled transcription/translation system and used this protein as prey in GST-RAB pull-down assays in which MyoE was detected with a rabbit polyclonal antiserum raised against its GTD. In these assays Sec4-GST and RAB11-GST, but not RAB5b-GST [RAB5b is the main EE RAB (Abenza et al., 2010)] pulled-down MyoE, showing discreet yet reproducible nucleotide specificity (Figure 1G). Nucleotide specificity was nevertheless established by experiments showing that myosin-5 was specifically retained by GST-RAB11 GTP*γ*S-affinity beads (Figure 2A, see below). Thus, Sec4 and RAB11 bind MyoE directly, focusing SVs at the SPK.

**Figure 2:**
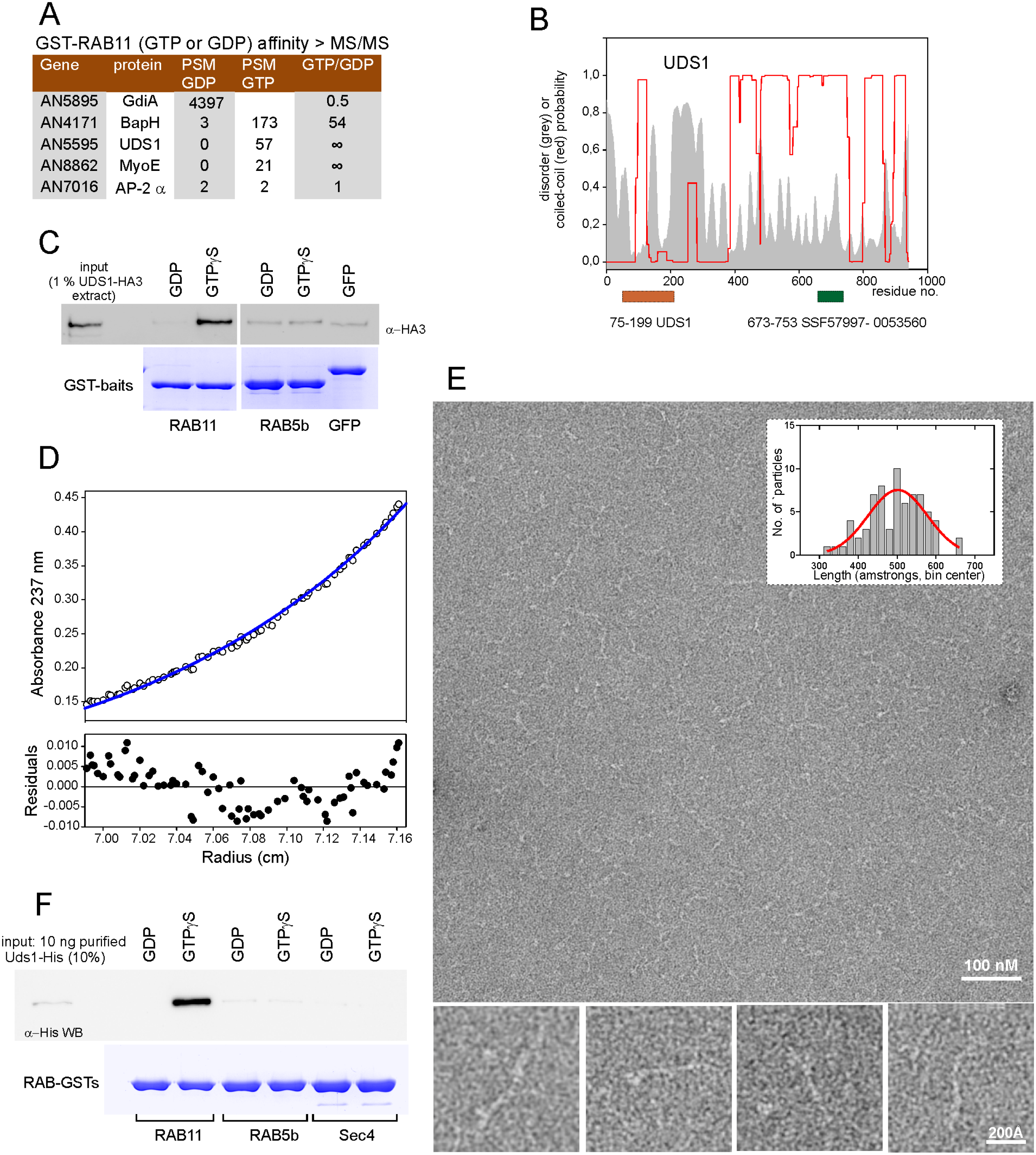
UDS1, a novel, direct effector of RAB11. (A) Table: Proteins retained by GTP*γ*S- and GDP-loaded RAB11-GST columns were analyzed in a QExactive mass spectrometer. The table lists the spectral counts obtained for each indicated protein and condition, as well as the relative enrichment detected in one sample vs. the other. Note that markedly abundant GdiA (GDP dissociation factor) interacts preferentially with GDP-RAB11. AP-2*α* was used as negative control. Bottom, GST-pull down assays with the indicated baits, using a prey extract of *A. nidulans* expressing UDS1-HA3 from the endogenously tagged gene. Pull-downs were analyzed by western blot with *α*-HA3 antibody. GST-GFP was used as a further negative control. (B) Features of UDS1. The probability of forming coiled-coils (red graph) and disordered regions (grey area) is shown. The positions of the UDS1 and SCOP superfamily domains are indicated. (C) UDS1 is a dimer *in vitro*. Equilibrium ultracentrifugation of bacterially expressed and purified UDS1 at a concentration of 4 µM; Top: The concentration gradient obtained (empty circles) is shown together with the best-fit analysis assuming that the protein is a dimer. Bottom plot, differences between experimental data and estimated values for the dimer model (residuals). (D) Negative-stain electron microscopy of purified UDS1. The proteins were stained with uranyl acetate and examined in a JEOL-1230 electron microscope. Four examples selected showed the extended screw-like form of UDS1. The lengths of *N=* 71 molecules were measured (plot; average 496 Å +/-73 S.D.). (E) GST-pull down assays using the indicated RABs and purified UDS1-His as prey. UDS1 was detected by anti-His western blotting.

### The actomyosin pathway protein UDS1 is a prototypic RAB11 effector

We hypothesized that other RAB11 effectors might reinforce the binding of MyoE to GTP-RAB11, similar to the situation with mammalian RAB11 and myosin-Vb (Hales et al., 2002, Schafer et al., 2014). To search for these effectors, we identified by liquid chromatography and tandem mass spectrometry (LC-MS/MS) the proteins retained by glutathione Sepharose beads containing RAB11-GST baits loaded with GDP or GTP*γ*-S. The resulting hits were ordered by abundance of peptide spectra matches (PSMs) in the GTP*γ*S sample relative to the GDP one, which helped to identify potential physiological hits. The highly abundant GDP-dissociation inhibitor GdiA (Pinar et al., 2015) served as specific GDP-RAB binder control, the previously characterized and abundant RAB11-GTP effector BapH (Pinar & Peñalva, 2017) served as positive control, and the unrelated AP-2 alpha-adaptin as negative one (Figure 2A). This analysis highlighted two potential actin-related hits. One was MyoE itself, which was exclusively retained by GST-RAB11 (GTP) beads, reinforcing the conclusion that this myosin-5 is a RAB11 effector. The second was the relatively abundant and highly specific RAB11-GTP effector AN5595 (Figure 2A). The 941 residue AN5595 product is predicted to have a strong tendency to form coiled-coils (Figure 2B). A *N. crassa* homologue of AN5595 denoted JANUS-1 interacts with the polarisome component Spa2 and has been suggested to serve as an SPK scaffold (Zheng et al., 2020). However, AN5595 showed features of an actomyosin regulator (Figure 2B), as it contains a SCOP Superfamily tropomyosin domain (SSF57997) suggestive of a parallel coiled-coil quaternary structure, and a UDS1 domain (PF15456) named after its as yet uncharacterized *Schizosaccharomyces pombe* homologue, whose name stands for ‘upregulated during septation’, and which localizes to the contractile actin ring in the mitotic septum (Ikebe et al., 2011). Therefore, we denoted AN5595 as UDS1.

To confirm that UDS1 is a *bona fide* RAB11 effector we HA3-tagged the protein endogenously and used USD1-HA3 cell extracts in pull-down assays with purified GST-RAB baits, loaded with GTP*γ*S or GDP, and with GST-GFP as negative control. UDS1-HA3 was pulled-down solely by GTP*γ*S-RAB11 but not by GFP, GDP-RAB11, GTP*γ*S-RAB5b or GDP-RAB5b baits (Figure 2C), confirming that UDS1 is subordinated to RAB11.

Next, we purified UDS1-His6 from bacteria. By gel filtration chromatography UDS1 eluted at a position corresponding to > 600 kDa (Figure S1), suggesting homo-oligomerization and/or a 3D structure substantially deviating from the globular shape. Sedimentation equilibrium ultracentrifugation of purified UDS1 (Mr 106,857 Da) revealed a buoyant mass of 57002 ± 403 Da corresponding to a molar mass of 209,073 Da ± 1612 Da, matching the molecular weight of a dimer (Figure 2D). Moreover, although the flexibility observed at the level of individual particles precluded us from obtaining 2D averages, individual EM images revealed a rod-shaped structure highly suggestive of a highly elongated coiled-coiled dimer. Therefore, UDS1 is an elongated dimer, with an approximate length of ∼500 Å (Figure 2E).

As the above RAB pull-down experiments using cell extracts do not rule out the possibility that RAB11 and UDS1 interact by way of bridging protein(s), we used the His-tagged protein to repeat the GST-RAB pull-down assays. Figure 2F shows that purified UDS1 behaves as the protein present in *Aspergillus* extracts, being pulled-down by GTP*γ*S-RAB11 but not by GDP-RAB11, nor by the inactive or active forms of RAB5b and Sec4. In summary, UDS1 is a coiled-coil dimer that binds directly to the (GTP) active form of RAB11.

### *Aspergillus* UDS1 colocalizes with both myosin-5 and RAB11 SVs

In current models (Figure 1A), RAB11 SVs arrive at the tip using kinesin-1 and are further concentrated at the SPK by myosin-5. Figure 3A shows that in agreement with these models RAB11 SVs fill a region at the apex that extends slightly beyond the SPK, as defined by the strictly apical MyoE-GFP signal. In colocalization experiments with RAB11, UDS1 behaves like MyoE, being restricted to the SPK, whereas RAB11 shows a slightly broader distribution (Figure 3B)(Movie 3). This distribution of RAB11 SVs extending beyond the SPK resembles the distribution of vesicles with a diameter of 70-90 nm observed by EM at the tip region (Hohmann-Marriott et al., 2006), which suggests that these vesicles correspond to RAB11 SVs. The behavior of UDS1, akin to that of MyoE, is consistent with UDS1 being a MyoE associate. Indeed, as predicted, UDS1 strictly colocalizes with myosin-5 in still images (Figure 3C) and across time, as seen in Figure 3C kymograph derived from a movie covering > 15 min (Movie 4). Moreover, in the absence of myosin-5 resulting in redistribution of RAB11 vesicles from the SPK to the apical dome (Figure 1A) UDS1 strictly colocalizes with RAB11 SVs arriving at the apical dome by way of MTs (Figure 3D), agreeing with UDS1 being a RAB11 effector that is present in SVs, rather than a structural component of the SPK. Movie 5 shows how UDS1-GFP recurs in the apical dome of a *myoE*Δ tip, as would be expected if UDS1 SVs arrive through MT transport to the PM.

**Figure 3.**
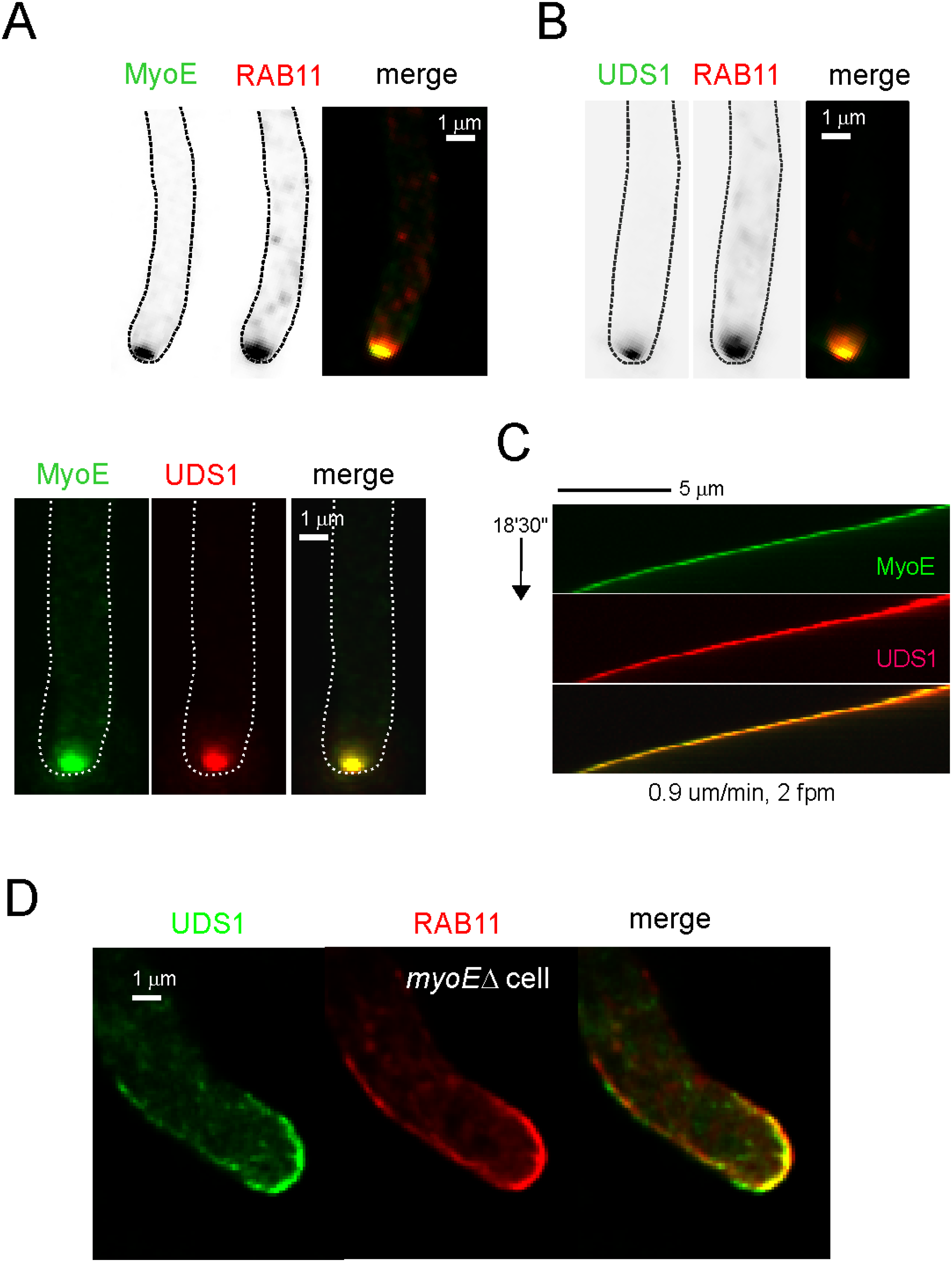
Subcellular localization of UDS1. (A) Representative images of a hyphal tip cell co-expressing mCh-RAB11 and MyoE-GFP. Images are MIPs of deconvolved Z-stacks. All images in this Figure are shown at the same magnification. (B) Images of a hyphal tip cell co-expressing mCh-RAB11 and endogenously tagged UDS1-GFP. Images are MIPs of deconvolved Z-stacks. (C) Left, images of a hyphal tip cell co-expressing endogenously tagged MyoE-GFP and UDS1-tdT. Images are MIPs of deconvolved Z-stacks. Right, MyoE-GFP and UDS1-tdT strictly colocalize across time: A 4D movie made with MIPs of Z-stacks acquired every 30 sec (Supplemental movie 3) was used to draw a kymograph across the long axis of a hypha growing at 0.9 µm/min. The diagonal lines traced by apical spots of the fluorescent proteins reflect the hypha growing by apical extension. (D) UDS1-GFP strictly colocalizes with mCh-RAB11 SVs transported by MTs to the apical dome in a cell lacking MyoE/myosin-5 (see also Figure 7A). Images are MIPs of deconvolved Z-stacks.

### Myosin-5 associates directly with HMSV, a fourth component of the RAB11/actomyosin pathway

To investigate the possibility that MyoE and UDS1 associate, we analyzed, by LC-MS/MS, GFP-Trap immunoprecipitates of MyoE-GFP and UDS1-GFP cells, using immunoprecipitates of a strain expressing the unrelated bait Uso1-GFP as a negative control (Uso1 acts in the ER/Golgi interface). UDS1 indeed pulled down MyoE, although MyoE pulled down UDS1 inefficiently, suggestive of weak or indirect interaction (Figure 4A). Remarkably, an as yet uncharacterized protein, the product of AN1213, coprecipitated with MyoE-GFP quite efficiently. Conversely, MyoE coprecipitated efficiently with GFP-tagged AN1213. A homologue of AN1213, denoted SPZ-1, has been investigated in *N. crassa* and proposed to serve as scaffold at the SPK (Zheng et al., 2020). However, for reasons that become clear below we denoted AN1213 as HMSV (*h*ooking *m*yosin to *SV*s). HMSV coprecipitated with UDS1-GFP as well, indicating that these proteins also interact (Figure 4A). In short, MyoE, UDS1 and HMSV appear to be associates, and components of the RAB11 pathway

**Figure 4.**
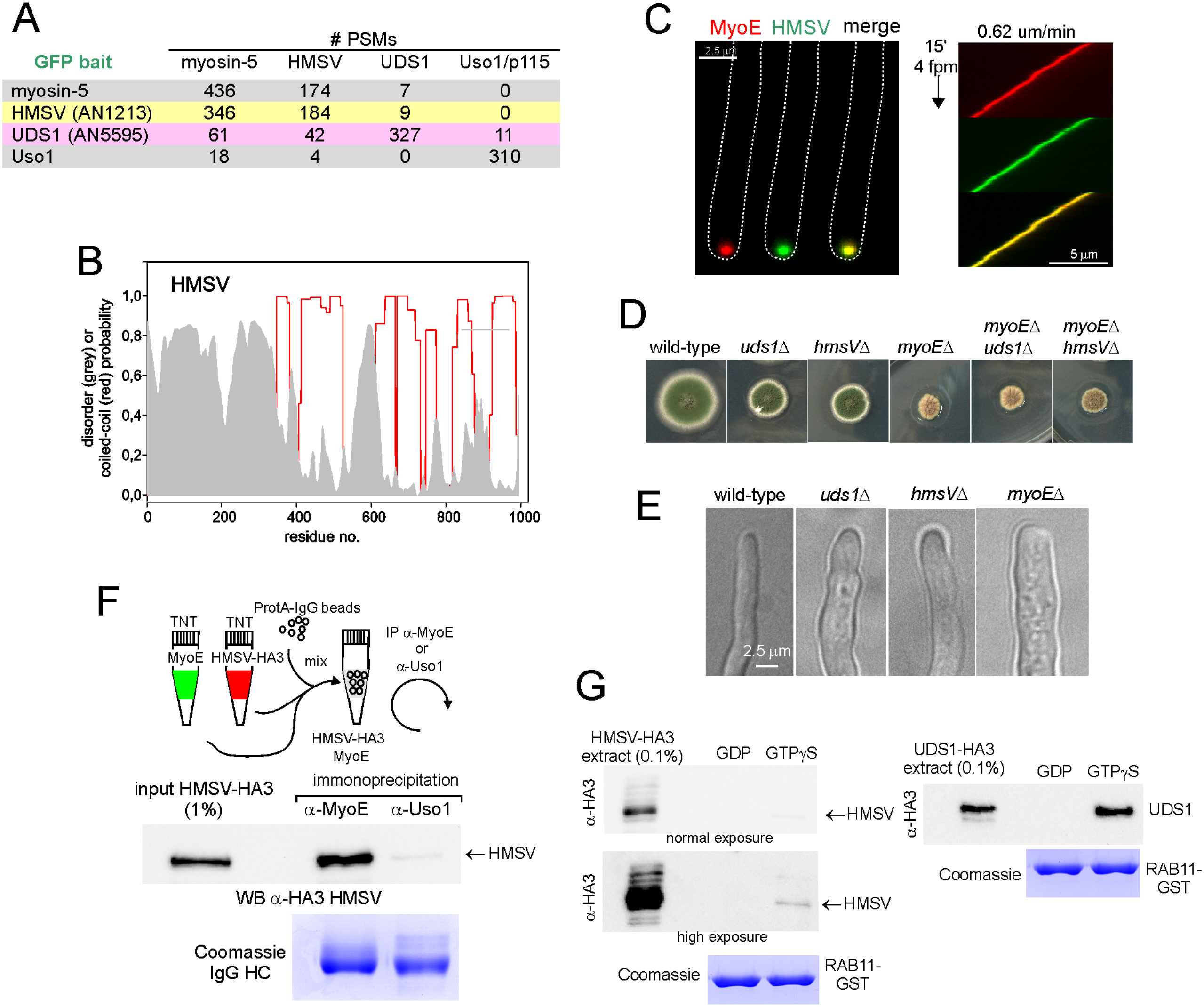
Characterization of HMSV, a myosin-5 interactor. (A) Cell extracts expressing the indicated GFP-tagged baits were immunoprecipitated with GFP-Trap and the proteins associated with them were analyzed by LC-MS/MS. The table lists the spectral counts obtained for each of the indicated co-precipitating proteins. The GFP-tagged unrelated protein Uso1 (the homologue of mammalian p115) was used as negative control. (B) Probability of forming coiled-coils (red graph) and disordered regions (grey area) along the primary sequence of HMSV. (C) Endogenously tagged MyoE-mCh and HMSV-GFP strictly colocalize. Left, green and red channel images are MIPs of deconvolved Z-stacks. Right, kymograph derived from Supplemental Movie 5, which is a 4D series constructed with MIPs of Z-stacks acquired with a beam splitter every 15 sec for a total of 15 min. The hypha was growing at 0.62 µm/min. (D) Growth tests at 37°C of strains with indicated genotypes, which were point-inoculated on complete medium. (E) Visual comparison of the different cell widths of the indicated mutant hyphae. (F) MyoE and HMSV interact directly: MyoE and HMSV-HA3 obtained from separate TNT reactions were combined with Protein A beads preloaded with polyclonal IgGs against MyoE or, as negative control, against the unrelated protein Uso1. Immunoprecipitates were analyzed by western blotting with *α*-HA antibody. Equal IgG load for the two IP reactions was confirmed by Coomassie staining of IgG’s heavy chains. (G) HMSV is not a (direct) RAB11 effector: GST-pull down assays with GDP and GTP*γ*S RAB11 baits, using prey cell extracts of *A. nidulans* expressing HMSV-HA3 (left) or, as positive control, UDS1-HA3 from endogenously tagged genes. Pull-downs were analyzed by western blot with *α*-HA3 antibody. Note that no signal of HMSV is detected under conditions in which the UDS1 signal is strong. However, after strong overexposure a faint band of HMSV is detectable in the GTP*γ*S reaction.

HMSV is a 994 residue-long protein whose 300 N-terminal residues are predicted to be disordered, while the remaining *∼*700 residues have strong propensity to form coiled-coils (Figure 4B). Like UDS1, HMSV localizes to the SPK, strictly colocalizing with MyoE (Figure 4C)(Movie 6). To determine the phenotypic consequences of removing UDS1 and HMSV we constructed null *uds1*Δ and *hmsV*Δ alleles. They are phenotypically indistinguishable, resulting in a radial colony growth defect (Figure 4D) and, at the cellular level, in abnormally wide hyphae (Figure 4E), both phenotypic features indicative of defective exocytosis. Notably, the colony growth defect resulting from *uds1*Δ and *hmsV*Δ was markedly weaker than that caused by *myoE*Δ (Figure 4D). Double *uds1Δ hmsV*Δ mutants behaved like the parental single mutants, consistent with the corresponding products being components of a functional unit (Figure S2). The fact that both *uds1*Δ and *hmsV*Δ are hypostatic to *myoE*Δ (Figure 4D) strongly suggested that this hypothetical complex acts through MyoE, although UDS1 and HMSV would not play an essential role in MyoE function.

The high yields of HMSV and MyoE recovered with their respective GFP-Traps immunoprecipitates (Figure 4A) strongly suggested that MyoE and HMSV are direct interactors. This prediction was confirmed by co-immunoprecipitation experiments using MyoE and HMSV-HA3 expressed by coupled transcription-translation reactions primed with their respective cDNAs. The two proteins were combined and immunoprecipitated with *α*-MyoE-specific IgGs or with IgGs raised against the unrelated protein Uso1 (acting in the ER/Golgi interface). *α*-MyoE IgGs, but not *α*-Uso1 IgGs, immunoprecipitated HMSV-HA3 (Figure 4F), establishing that HMSV and MyoE interact directly. In short, all the above data strongly suggested that HMSV acts as a connector between RAB11/UDS1 and MyoE.

Unlike UDS1, HMSV did not appear to interact with RAB11 in shotgun proteomic experiments (Figure 2A). To confirm this observation, we performed more sensitive GST-pull down assays with extracts of cells expressing HA3-tagged baits. Under conditions in which UDS1-HA3 strongly associated with RAB11-GST, HMSV-HA3 did not (Figure 4G). Even though strong overexposure of the blots revealed a faint signal in the GTP*γ*S-RAB11 lane, these data argued against HMSV being a direct interactor of RAB11 and suggested instead that another factor might bridge HMSV to RAB11 (note that total cell extracts *—*not purified proteins*—* were used as preys in this experiment).

### HMSV scaffolds a myosin-5-containing heterotrimeric complex that binds to the active RAB11 conformer

An appealing candidate to link HMSV indirectly to RAB11 was UDS1. Figure 5A shows that GST-UDS1, but not the unrelated bait GST-GFP, pulled-down *in vitro* synthesized HMSV-HA3. In contrast, neither GST bait pulled-down *in vitro* synthesized Uso1-HA3, confirming specificity and establishing that purified UDS1 and HMSV interact. Therefore, by interacting directly with both MyoE and UDS1, HMSV would act as scaffold of a heterotrimeric complex that is recruited by RAB11 to SVs by contacting both UDS1 and MyoE/myosin-5.

**Figure 5.**
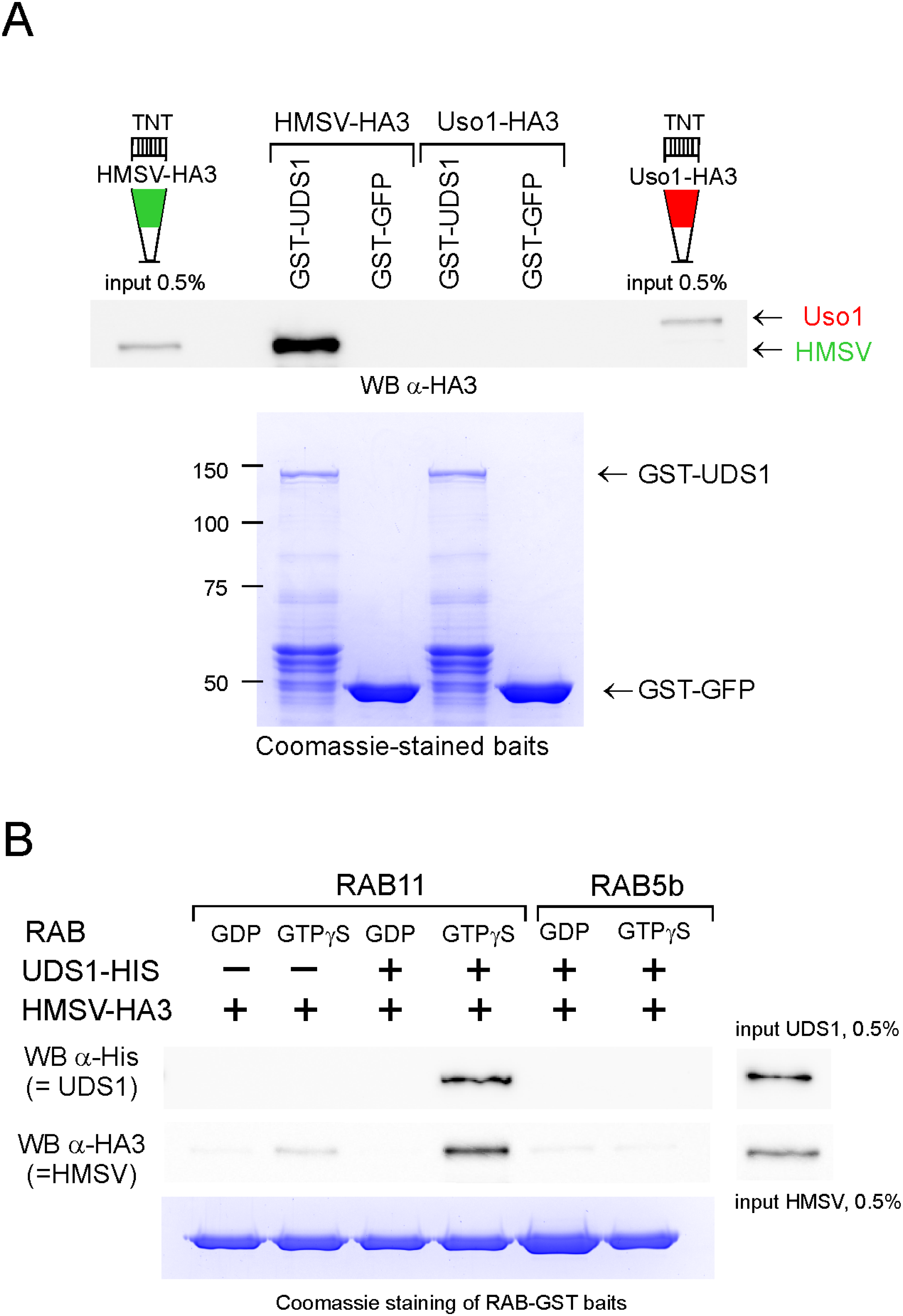
HMSV is a direct interactor of UDS1 and an indirect effector of RAB11. (A) HMSV and UDS1 interact directly: Pull-down assays with GST-UDS1 as bait and HMSV-HA3 or, as negative control, Uso1-HA3 as preys. Preys were obtained by TNT expression. Pull-downs were analyzed by *α*-HA3 western blotting. (B) UDS1 bridges the interaction of HMSV with the active form of RAB11. GST pull-down assays with RAB11 and, as negative control, RAB5b baits. The preys, combined in the reactions as indicated, were purified UDS1-His and TNT-synthesized HMSV-HA3. Samples of the reactions were analyzed by *α*-His and *α*-HA western blotting. HMSV is recruited to RAB11 only when UDS1 is present and only if the GTPase is in the active conformation.

To test this model, we performed two sets of experiments. First, we demonstrated *in vitro* that HMSV is recruited to active RAB11 only if UDS1 is present to bridge the interaction (Figure 5B). We performed GST-RAB pull-downs in the presence of bacterially expressed UDS1, *in vitro* synthesized HMSV-HA3 or both. HMSV was recruited by GTP*γ*S-RAB11, but did so only when UDS1 was present in the reaction mix. Neither conformation of RAB5b nor GDP-RAB11 pulled-down HMSV even when UDS1 was present. We conclude that HMSV is an indirect effector of RAB11 that requires the presence of UDS1 to be recruited to the GTPase.

Secondly, we demonstrated that the stable complex reconstructed *in vitro*, consisting of MyoE, HMSV and UDS1, is present in cellular lysates and is scaffolded by HMSV. As determined by anti-MyoE Western blotting of GFP-Trap immunoprecipitates of whole-cell extracts, MyoE strongly associates with UDS1-GFP and with HMSV-GFP, but not with the unrelated bait Uso1-GFP (Figure 6A). Indeed, the associations are so efficient that co-immunoprecipitated MyoE could be visualized directly by silver-staining of SDS-PAGE gels (Figure 6A, right). Despite HMSV appears to be the less abundant bait (anti-GFP western blot, Figure 6A, right), the interaction between MyoE and HMSV was markedly more efficient than that between MyoE and UDS1, in agreement with the fact that MyoE and UDS1 interact indirectly by way of HMSV. Consistently, the interaction between MyoE and UDS1 was undetectable with *hmsV*Δ extracts (i.e. was completely dependent on the presence of HMSV) (Figure 6B), whereas than that between MyoE and HMSV was completely independent of UDS1, taking place irrespectively of whether wild-type or *uds1*Δ extracts were used (Figure 6C). Lastly, the interaction between UDS1-GFP and HMSV-HA3 was completely independent of MyoE (Figure 6D), as predicted by *in vitro* reconstitution experiments above.

**Figure 6.**
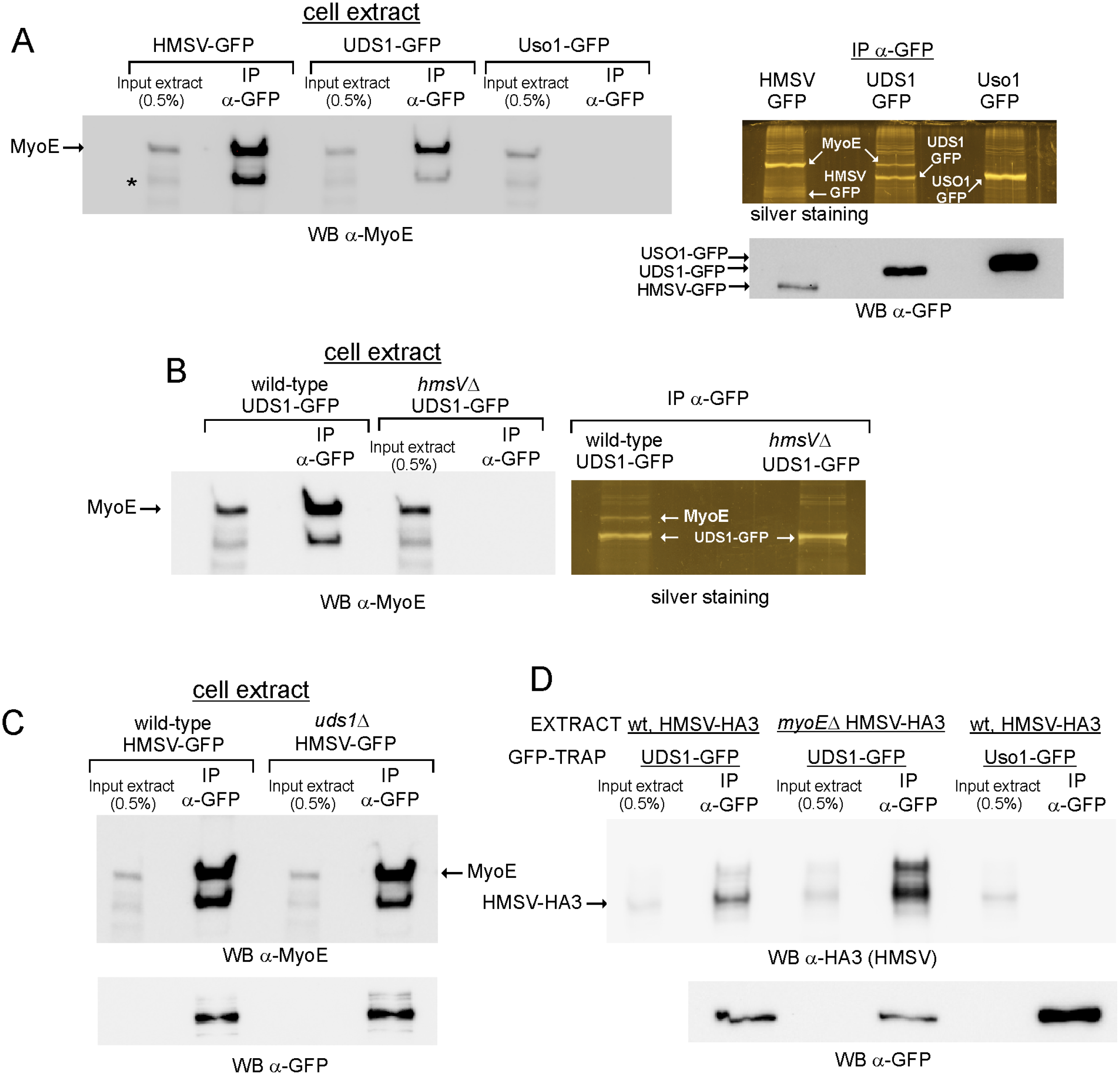
In cells, UDS1 and MyoE/myosin-5 are present in a complex scaffolded by HMSV. (A) Myosin-5/MyoE associates with UDS1 and HMSV but not with Uso1. Extracts of cells expressing indicated endogenously-tagged GFP proteins were immunoprecipitated with GFP-Trap nanobody. Left, western blot analysis of immunoprecipitates with *α*-MyoE antibody. The band indicated with an asterisk is unspecific. Right top, silver staining of immunoprecipitates. MyoE is remarkably abundant in the HMSV sample, detectable in the UDS1 sample and absent in the Uso1 sample. Right bottom, relative levels of the preys revealed by *α*-GFP western blotting. (B) UDS1 and MyoE/myosin-5 associate only if HMSV is present. (C) HMSV and MyoE/myosin-5 associate efficiently when UDS1 is absent (D) UDS1 associates with HMSV regardless of whether MyoE is absent or present. GFP-Trap immunoprecipates of cells co-expressing HMSV-HA3 were analyzed by *α*-HA3 western blotting to detect HMSV, and by *α*-GFP western blotting to reveal the relative levels of the baits.

Taken together these data show that these proteins form a complex in the order MyoE/HMSV/UDS1 that has the dual ability to interact with the active form of RAB11 through UDS1-(Figures 3-5) and MyoE-mediated (Figure 1) contacts.

### Evidence that UDS1 and HMSV are a co-receptor assisting RAB11 to recruit myosin-5 to SVs

A diagnostic readout of myosin-5 transport is the focusing of SVs at the SPK. Consistent with UDS1 and HMSV acting in a complex regulating actomyosin transport, both *uds1*Δ and *hmsV*Δ affected RAB11 SVs similarly, reallocating them from the SPK to a crescent-shaped distribution in the apical dome (Figure 7A). This effect was markedly less prominent than that caused by *myoEΔ,* which resulted in a broader crescent and, as discussed above, in a marked reduction of the signal of SVs docked at the tip cortex (Figure 1B). Therefore, these data strongly indicate that myosin-5 transport is debilitated in *uds1*Δ and *hmsV*Δ mutants, such that although this transport is not abolished, MT/kinesin-1-mediated transport gains prominence, which results in targeting SVs to a broader surface determined by the sites at which MTs’ plus ends reach the apical dome.

**Figure 7.**
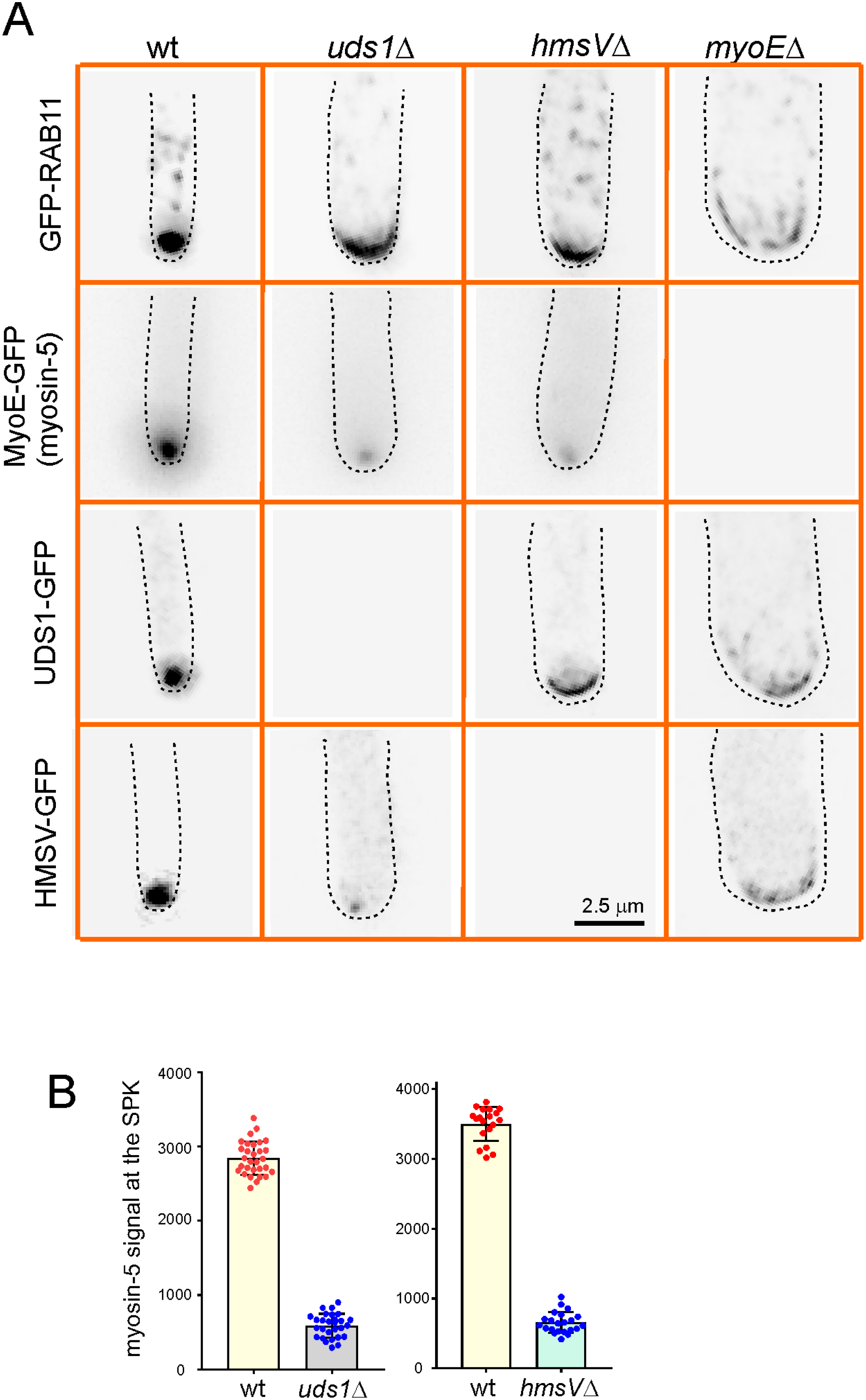
Localization of myosin-5 and of RAB11 and its associates in different genetic backgrounds supports the role of the tripartite complex as myosin-5 adaptor. (A) Localization of the components of the tripartite complex in the different genetic backgrounds. Images are shown at the same magnification. All GFP reporters were endogenously tagged, implying that the corresponding null background images are empty. Images are MIPs of deconvolved Z-stacks. (B) Quantitation of the MyoE-GFP signal in the SPK of *uds1*Δ and *hmsV*Δ cells compared to the wt. Mean values for the left graph were: wild-type 2842 A.U. ± 227 S.D. *vs. uds1*Δ 558 ± 159 S.D., *P* < 0.0001 in unpaired *t-*test. For the right graph (corresponding to a different experiment), wild-type, 3496 ± 245 S.D. *vs.* 654 ± 149 in the *hmvS*Δ mutant (*P* < 0.0001 in unpaired *t*-test).

Impairment of actomyosin transport in these mutants explains the partial exocytic deficit that growth tests indicate (Figure 4D).

All the above experiments suggested that the UDS1/HMSV complex might form part of a co-receptor reinforcing the RAB11-mediated recruitment of myosin-5 to SVs. Myosin-5 dwells in an inactive conformation that is shifted to the active conformation by cargo (Donovan & Bretscher, 2015). Thus, a deficit in cargo loading would be translated into a drop in myosin-5 activity, which should in turn result in a reduction in the levels of myosin-5 at the SPK. Figure 7A, and B shows that both *uds1*Δ and *hmsV*Δ reduce the SPK MyoE signal by 5-6-fold (a significant difference; *P <* 0.0001 in unpaired *t-*tests), supporting the contention that in these mutant backgrounds the loading of myosin-5 with SVs is compromised. Movie 7 dynamically depicts the effect of *hmsV*Δ on the steady-state levels of MyoE accumulating at the SPK.

We next investigated the dependence of UDS1 and HMSV localization on each other. In *hmsV*Δ cells UDS1 delocalized from the SPK to an apical crescent remarkably similar to that observed with RAB11 (Figure 7A), indicating that the connection of UDS1 (RAB11) SVs with MyoE is impaired (recall that a broader distribution indicates that the balance between actomyosin and MT transport has been shifted towards the latter). In sheer contrast, HMSV is not delocalized from the SPK in *uds1*Δ cells, but the signal was reduced to an extent roughly commensurate with the reduction in MyoE signal (Figure 7A and B), indicating that HMSV goes with the proportion of myosin-5 that is (less efficiently) loaded with cargo by way of the direct interaction of RAB11 with the motor, and therefore that HMSV binds indirectly to RAB11 by way of UDS1. Notably, the localization of RAB11, UDS1 and HMSV in *myoE*Δ cells is remarkably similar (Figure 7A) (Movie 8 for HMSV), reflecting their dependence on MT transport in their distribution to the apical dome, and further demonstrating that UDS1 and HMSV are components of RAB11 SVs rather that structural constituents of the SPK.

### Ablation of UDS1 or HMSV impairs the delivery of an exocytic cargo to the SPK

A well characterized cargo of RAB11 SVs is the chitin synthase ChsB (Hernández-González et al., 2018a). This integral membrane protein is exocytosed to the apical plasma membrane by way of the SPK, diffuses away from the tip and it is taken up by a highly active endocytic collar that transports it to a sorting endosome. From this compartment ChsB returns to the TGN where it is incorporated into RAB11 SVs delivered to the SPK (Figure 8, scheme). In the wild-type, a proportion of ChsB is present in the SPK. In *uds1*Δ and *hmsV*Δ cells this accumulation of ChsB in the SPK is no longer seen, resembling the situation with RAB11, which is included in Figure 8 for comparison. We interpret that the absence of the UDS1/HMSV co-receptor affects transport of a RAB11 cargo from the TGN to the SPK; (In passing, reduced delivery of ChsB to the apex might contribute to the morphological defect characteristic of *uds1*Δ and *hmsV*Δ hyphae).

**Figure 8.**
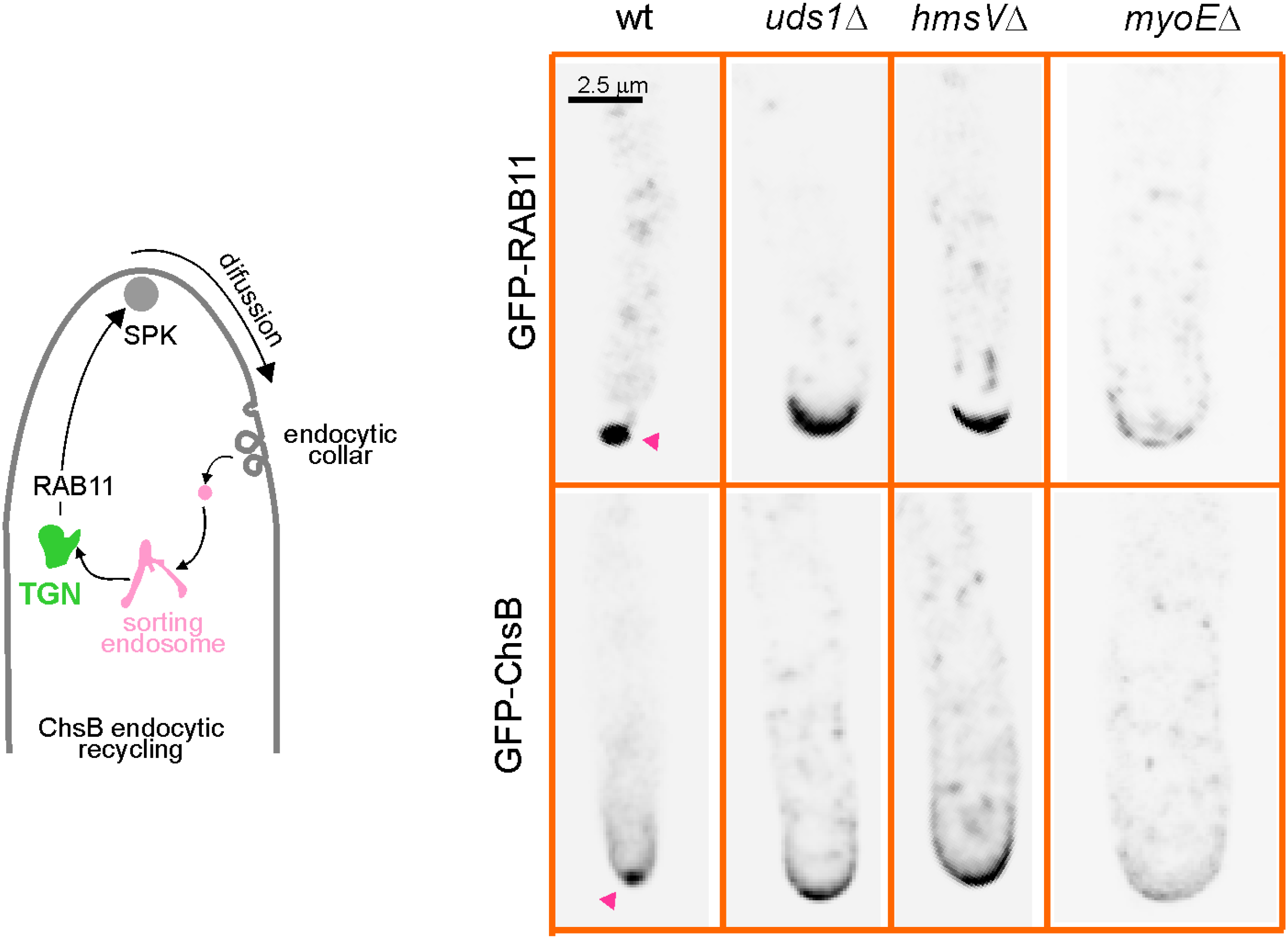
A cargo of the RAB11 recycling pathway is delocalized from the SPK in *uds1*Δ and *hmsV*Δ cells. The scheme depicts the endocytic recycling pathway followed by the chitin-synthase ChsB. *uds1*Δ and *hmsV*Δ mutations delocalize ChsB from the SPK, correlating with delocalization of RAB11 from the SPK to an apical crescent. Red arrowheads indicate the SPKs.

In summary, our data strongly support a model in which the HMSV and UDS1, which are indirect and direct RAB11 effectors, respectively, serve as co-adaptor between SVs budding from the TGN and MyoE, the only *Aspergillus* myosin-5 motor (Figure 9). When this co-receptor is disorganized by the ablation of either of its two components, actomyosin transport of these SVs still occurs, albeit less efficiently, due to the direct interaction between RAB11 and MyoE.

**Figure 9.**
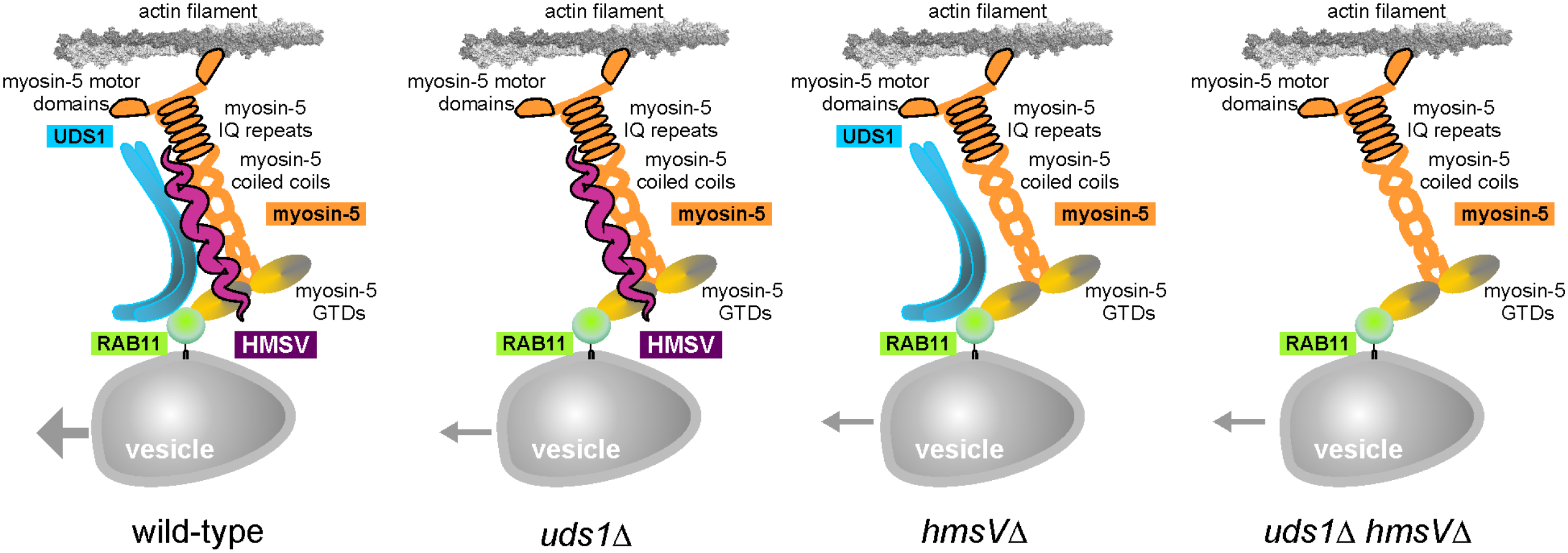
A model for the engagement of myosin-5 by RAB11 SVs in *A. nidulans*. In the wild-type RAB11 is recruited to SVs during the Golgi-to-post-Golgi transition. RAB11 interacts with the GTD of MyoE and with the UDS1 dimer. UDS1 in turn connects active RAB11 to the HMSV scaffold (represented here as a monomer, but potentially being a dimer). HMSV bridges RAB11/UDS1 to the MyoE myosin-5. In the presence of the whole complex myosin-5 transport is most efficient (large arrow). In the absence of UDS1 or HMSV there is still myosin-5 transport due to the direct interaction between RAB11 and the MyoE GTD, albeit this transport is markedly less efficient (small arrows), such that the accumulation of SVs in the SPK is impaired and MT-dependent transport becomes more prominent, leading to the characteristic apical dome distribution of SVs in these mutants (Figure 7A). Phenotypically the *hmsVΔ uds1*Δ double mutant is indistinguishable from either of the single mutant strains (Figure S2). HMSV and UDS1 might sustain efficient myosin-5 transport by reinforcing the interaction between RAB11 and the motor or, alternatively, they might increase processivity of the motor or facilitate the switch from MTs to actin cables in the crowded cytoplasm of the hyphal tip.

## Discussion

The ability of myosin-5 to transport cargo along actin filaments is crucial for the biogenesis and distribution of membranous compartments (Hammer & Sellers, 2012, Wong & Weisman, 2021). *A. nidulans* has a single myosin-5, MyoE (Taheri-Talesh et al., 2012), implying that specificity for different cargoes must be mediated by different adaptors, *i.e.* proteins engaging membranous organelles to the motor (Cross & Dodding, 2019, Wong & Weisman, 2021). Adaptors often involve a RAB family member, exploiting the fact that individual RABs display a high selectivity for a cognate membrane compartment (Pfeffer, 2013, Pinar & Peñalva, 2021). RABs can interact directly with myosin-5, or indirectly, by means of intermediate proteins that bridge the activated RAB and the motor (Hammer & Sellers, 2012, Wong & Weisman, 2021). A well understood example of co-adaptor is melanophilin, which bridges RAB27 on melanosomes to MyoVa (Wu et al., 2002). Even if the binding of the RAB to the myosin-5 is direct, it frequently involves additional co-adaptors that help stabilizing the complex. This is the case of metazoan RAB11, FIP2 and MyoVb (FIP2 is a direct effector of RAB11), which form a tripartite complex required for traffic between recycling endosomes and the PM (Hales et al., 2002, Li et al., 2007, Schafer et al., 2014, Wang et al., 2008). Besides complex stabilization co-adaptors play additional roles. For example, the C-terminal region of melanophilin binds F-actin, and this binding dramatically increases the processivity of MyoVa (Sckolnick et al., 2013). Moreover, melanophilin tracks the plus-ends of MTs by hitchhiking on EB1. In turn, melanophilin recruits MyoVa to the MTs’ plus ends, which might be mechanistically important to ensure the efficient transfer of melanosomes from MTs to actin cables.(Wu et al., 2005). Another example of additional functions of RAB-scaffolded myosin-5 adaptors occurs in mouse oocytes, where myosin-5 is recruited to RAB11 vesicles by cooperative interactions with both the GTPase and the actin nucleator SPIR-2, which helps coordinating MyoVa vesicle transport with actin nucleation (Pylypenko et al., 2016). Adaptors may play additional roles unrelated to transport itself. Notably, they may help to release the vesicle from myosin-5 upon arrival to destination: Phosphorylation and ubiquitin-mediated degradation of the yeast vacuolar adaptor Vps17 is required to release the organelle from Myo2 (Wong et al., 2020). We cannot rule out that UDS1 and HMSV play additional roles, such as retaining SVs in the SPK/vesicle supply center. For example, their *N. crassa* homologues, which are not involved in vesicle trafficking, have been proposed to act as polarity scaffolds (Zheng et al., 2020).

In *A. nidulans* the biogenesis of SVs dispatched to the PM is mediated by RAB11. SVs are loaded with myosin-5, kinesin-1 and dynein (Pantazopoulou et al., 2014, Peñalva et al., 2017), but the adaptors linking these molecular motors to vesicles remain uncharacterized. Resembling melanosome transport, kinesin-1 hauls SVs to tip-proximal regions before transferring them to myosin-5, which concentrates SVs at the SPK (the cell periphery, in the case of melanosomes). This two-step mechanism involves the transfer of SVs from MT- to F-actin-mediated transport, a relay that is almost certainly compromised by the high density of cytoskeletal tracks and organelles populating the hyphal tip. Genetic and biochemical evidence showed that RAB11 engages MyoE (= myosin-5) by way of a direct contact between the GTPase and the thoroughly studied GTD of the motor (GTD cargo binding domains of mammalian MyoVa,b,c and fungal Myo2 and MyoE are conserved (Pashkova et al., 2006, Pylypenko et al., 2013)). Yet we hypothesized that additional co-receptors were likely to be involved to facilitate an efficient relay and the subsequent MyoE-powered journey of SVs to the SPK across a crowded cytoplasm, and that these would be effectors of RAB11. Shotgun proteomics identified UDS1 and HMSV as components of a tripartite complex (see the model in Figure 9), in which at least one component, UDS1, is dimeric. UDS1 binds directly to both RAB11 and HMSV, and HMSV is a scaffold that binds directly to both MyoE and UDS1, connecting the motor to RAB11-UDS1. In agreement, in *hmsV*Δ cells UDS1-GFP distributes like RAB11 whereas in *uds1*Δ cells HMSV-GFP distributes like myosin-5, indicating that the whole receptor can be split in two stable subcomplexes (Figure 7). Therefore, both UDS1 and HMSV are necessary for the assembly of a receptor complex whose absence results in debilitated F-actin-mediated transport, which is reflected in the spreading of RAB11 SVs across the apical dome. Inefficient F-actin transport of RAB11 correlates with slower colony growth resulting from ablation of either co-adaptor, and spreading of RAB11 SVs across the hyphal tip dome correlates with delocalization of its cargo, ChsB, from the SPK. Of note, F-actin transport is not abolished without the co-adaptors (Figure 9), because RAB11 is able to bind MyoE directly, which makes the phenotypic consequences of ablating UDS1 or HMSV less deleterious than those resulting from removing MyoE, whose ablation leaves SV transport exclusively in the hands of kinesin-1.

As to how the presence of two additional proteins contributes to the efficiency of myosin-5 transport it is worth mentioning that in the absence of MyoE a proportion of RAB11 SVs decorates the array of tip actin cables emanating from the SPK (Pantazopoulou et al., 2014), suggesting that these vesicles contain a F-actin-binder. It is tempting to speculate that MyoE co-adaptor(s) resemble melanophilin (Sckolnick et al., 2013) or the Sec4p::Myo2p co-adaptor Smy1p (Hodges et al., 2009, Lwin et al., 2016) in that they interact with actin cables to increase the processivity of the motor. Even more suggestive is the hypothetical possibility that F-actin binding by the MyoE co-adaptors facilitates the switch between MT and F-actin transport. It should be noted that Sec4 cannot recruit UDS1 (Figure 2E), which is consistent with the view that *A. nidulans* Sec4 acts downstream of the RAB11-mediated transport of SVs to the SPK, mediating the ultimate step of exocytosis (Pinar & Peñalva, 2021).

Our work highlights the fact that cargo adaptor proteins for myosin-5 are difficult to identify by primary sequence or domain composition-based searches (Wong & Weisman, 2021). Both UDS1 and HMSV are predicted coiled-coil proteins. Cross and Dodding have recently reviewed the frequent occurrence of coiled-coil proteins among adaptors of molecular motors, including myosin-5 (Cross & Dodding, 2019). A well-understood example is the coiled-coil melanosome protein RILPL2 (RAB interacting lysosomal protein-like 2), which bridges RAB36 with the MyoVa GTD (Matsui et al., 2012, Wei et al., 2013).

In summary, we have identified a novel receptor complex required for the efficient coupling of RAB11 SVs to the myosin-5 MyoE. Proof-of-concept that a motor-cargo interface can be targeted by a small chemical has been recently provided (Randall et al., 2017). Although speculative at the moment, the possibility of interfering with fungal growth by diminishing the efficiency of myosin-5 mediated transport is appealing.

## Methods

### Aspergillus techniques

Standard *A. nidulans* media were used for strain propagation and conidiospore production. GFP and epitope-tagged alleles were introduced in the different genetic backgrounds by meiotic recombination (Todd et al., 2007) and/or transformation (Tilburn et al., 1983), which used recipient *nkuA*Δ strains deficient in the non-homologous end joining pathway (Nayak et al., 2005). Complete strain genotypes are listed in Table S1.

### Null mutant strains and protein tagging

*uds1*Δ, *hmsV*Δ, *sec4*Δ (Pantazopoulou et al., 2014) and *myoE*Δ (Taheri-Talesh et al., 2012) were constructed by transformation-mediated gene replacement with cassettes made by fusion PCR carrying appropriate selectable markers (Szewczyk et al., 2006). Integration events were confirmed by PCR with external primers.

The following proteins were C-terminally tagged endogenously, using cassettes constructed by fusion PCR (Nayak et al., 2005, Szewczyk et al., 2006): UDS1-GFP, UDS1-HA3, UDS1-tdTomato, HMSV-GFP, HMSV-HA3, MyoE-GFP (Taheri-Talesh et al., 2012), MyoE-mCherry and ChsB-GFP (Hernández-González et al., 2018a). GFP-RAB11 (Pantazopoulou et al., 2014) and mCherry-RAB11(Pinar & Peñalva, 2020) were expressed from its own promoter.

### GFP-GTD transgene

A transforming cassette consisting of, from 5’ to 3’, the sucrose-inducible promoter of the inulinase gene (*inuA*) (Hernández-González et al., 2018b), the GFP-coding sequence translationally fused to the coding sequence for residues 1082 through 1569 of MyoE, the *A. fumigatus pyrG* gene and the *inuA* gene 3’-flanking region was constructed by 5-way fusion PCR, using the following primers (underlined sequences indicate regions of overlap used for fusion PCRs):

(1): *inuA* promoter region, PCR-amplified with primers: 5’-GTGGAGGCCACTCTCGGAAAC-3’ 5’-<u>CAGTGAAAAGTTCTTCTCCTTTACTCAT</u>TTTGGTGATGTCGCTGACCGC-3’ (the underlined overlapped with GFP-coding region)
(2): GFP-(Gly-Ala)_6_ 5’- ATGAGTAAAGGAGAAGAACTTTTC-3’ 5’- GGCACCGGCTCCAGCGCCTGC-3’
(3): MyoE-GTD 5’- <u>CTGGTGCAGGCGCTGGAGCCGGTGCC</u>CAGGCGTTGAACGGAGACCAGC- 3’: [the underlined overlapping with the GFP-(Gly-Ala)_6_] coding region. 5’- <u>ATTCCAGCACACTGGCGGCCGTTAC</u>TTACTCCATCACCCCATTCTCAG-3’: (the underlined overlapping with *pyrGAf*)
(4): *pyrGAf* 5’- GTAACGGCCGCCAGTGTGCTG-3’ 5’- GTCTGAGAGGAGGCACTGATG-3’
(5): *inuA* 3’-UTR 5’- <u>ACGCATCAGTGCCTCCTCTCAGAC</u>AGGATCTAGCTAGATGTTTTGTTG-3’: 5’- CAGCAGTCAAGCAATACCAAGC-3’ (the underlined overlapping with *pyrGAf*).

The cassette was used to replace the *inuA* gene, considered to be a safe haven, by homologous recombination. *inuA*Δ does not affect growth on carbon sources other than inulin (glucose is used as standard carbon source in *A. nidulans* media).

### Myosin-5 mutants

A DNA cassette consisting of the 1478 3’-coding nucleotides of *myoE*, the *A. fumigatus pyrGAf* gene and 870 nucleotides of the *myoE* 3’-UTR region was cloned in pGEM-T easy (Promega). This plasmid was used as template for site-directed mutagenesis (QuickChange kit, Agilent Technologies) with the following oligonucleotides:

(a) Y1414R (genomic sequence coordinates T4391C A4392G): Fw: 5’- GAGGCCTCCAGATCAACCGCAACATAACTCGCATCGAG-3’ Rv: 5’- CTCGATGCGAGTTATGTTGCGGTTGATCTGGAGGCCTC-3’
(b) K1472E (A4658G): Fw: 5’- CTCTCCAAACCAAATCCAAGAGCTGCTAAACCAATACCT-3’ Rv: 5’- AGGTATTGGTTTAGCAGCTCTTGGATTTGGTTTGGAGAG-3’

Wild-type and mutant DNA cassettes, obtained after *NotI* digestion, were used to transform strain MAD5736. Transformants incorporating the wild-type and mutant *myoE* alleles were identified by PCR and confirmed by DNA sequencing. A transformant carrying the wild-type cassette was indistinguishable from the ‘true’ wild-type in terms of growth.

### Plasmids for protein expression

#### GST constructs

pET21b-RAB11-GST: carries cDNA encoding cysteine-less RAB11 with GST C-terminally attached. *Nde*I/*BamH*I insert in pET21b.

pET21b- Sec4-GST: carries cDNA encoding cysteine-less Sec4 with GST C-terminally attached. *Nde*I/*Xho*I insert in pET21b.

pET21b-RAB5b-GST: carries cDNA encoding cysteine-less RAB5b with GST C-terminally attached. *Nhe*I/*Not*I insert in pET21b.

Note that in all three constructs GST is attached to the C-termini of the RABs, and that they all include a stop codon after the GST coding region to interrupt translation before the His tag.

pGEX2T-GFP: pGEX-2T derivative encoding a GST-sGFP fusion.

pGEX2T-UDS1: UDS1 cDNA cloned as *BamH*I in pGEX-2T (N-terminal GST)

#### TNT expression constructs

pSP64 MyoE: MyoE cDNA cloned as a *BamH*I fragment in Promega’s pSP64 poly (A). pSP64 HMSV-HA: C-terminally HA3-tagged cDNA encoding HMSV cloned as *Pst*I/*Xma*I in Promega’s pSP64 poly (A).

pSP64 UsoA-HA: C-terminally HA3-tagged cDNA encoding Uso1 cloned as *Pst*I/*Xma*I in Promega’s pSP64 poly (A).

#### UDS1-His6 construct

pET21b-UDS1: UDS1 cDNA cloned as NheI/NotI in pET21b

### *In vitro* transcription/translation

Proteins were synthesized using TNT*®* SP6 Quick Coupled Transcription/Translation system (Promega L2080) using the standard reaction mix (rabbit reticulocyte lysate plus amino acids) supplemented with 20 µM methionine. Reactions were primed with 1 µg of purified, circular pSP64 derivatives, which were purified using NucleoBond Xtra-Midi columns (Macherey Nagel, FRG).

### Antibodies and western blotting

Antisera against MyoE and Uso1 were raised in rabbits by Davids Biotechnology (https://www.davids-bio.com). Animals were immunized with His6-tagged polypeptides containing residues 1082-1569 of MyoE (the GTD) or residues 1-659 of USO1. These polypeptides were purified by Ni2+ affinity chromatography after expression in *E. coli* BLB21 as described (Pinar et al., 2015). Antibodies against the target proteins were purified from raw antisera (40 ml) by affinity chromatography with the respective antigens, previously coupled to 1 ml HI-TRAP NHS columns (GE/Merck) packed with Sepharose pre-activated for covalent coupling of ligands containing primary amino groups, following instructions of the manufacturer. Antibodies were eluted with 100 mM glycine, pH 3.0, neutralized with 2M Tris, pH 7.5 and stored at −20°C.

Western blots were reacted with the following antibodies:

#### For HA3-tagged proteins

*α*-HA rat mAb (1/1,000) (Roche) as primary antibody, and HRP-conjugated *α*-rat IgM+IgG, as secondary antibodies (Southern Biotechnology; (1:4,000).

#### For His6-tagged UDS1

*α*-His primary antibody (1/10,000; Clontech) and HRP-conjugated goat anti-mouse IgG (H+L) secondary antibodies (Jackson Immunoresearch, 1/5000).

#### For MyoE

MyoE/myosin-5 was detected with a custom-made *α*-MyoE-GTD antiserum (1/4000; see above) and donkey HRP-coupled *α*-rabbit IgG (GE NA-934) as secondary antibodies.

#### For *α*-GFP western blotting

we used Roche’s mixture of two mouse mAbs (1/5000) as primary antibodies and HRP-conjugated AffiniPure goat anti-mouse IgG (H+L) secondary antibodies (Jackson Immunoresearch, 1/5000). In all cases reacting bands were detected with Clarity western ECL substrate (Biorad Laboratories).

### RAB-GST purification and nucleotide loading

500 ml bacterial cultures in LB plus antibiotics as appropriate were incubated at 37°C until reaching a O.D. of 0.6-0.8 at 600 nm. These primary cultures were induced with 0.1 mM IPTG, transferred to a 15°C incubator and shaked for an additional 20 h. Cells were collected by centrifugation and stored at −80°C. A pellet corresponding to 250 ml of the culture was resuspended in PBS containing Roche’s cOmplete^TM^ protease inhibitor cocktail, 0.2 mg/ml lysozyme and 1 µg/ml of DNAse I (Abenza et al., 2010) and lysed in a French Press. After centrifugation at 30,000 x g and 4°C for 30 min, the supernatant was mixed with 300 µl of glutathione-Sepharose 4B (Sigma) and incubated at 4°C for 1 h in a rotating wheel. Sepharose-bound RABs were resuspended in a buffer consisting of 25 mM HEPES PH 7.5, 110 mM KCl, 1 mM DTT, 10 mM EDTA and 125 µM GDP o GTPγS and incubated for 30 min at 30°C with gentle rocking. Beads were then washed twice with nucleotide loading buffer (as above, but containing 10 mM Cl2Mg instead of 10 mM EDTA) before incubating them overnight at 25°C with gentle rocking in nucleotide loading buffer containing GDP or GTPγS (Jena Bioscience UN-1172 and UN-412, respectively).

### UDS1-His6 expression and purification from bacteria

*E. coli* cells (BLB21 pRIL) carrying pET21b-UDS1 were cultured at 37°C in LB containing ampicillin and chloramphenicol until reaching and OD^660^ of 0.5. At this point cultures were induced with 0.1 M IPTG, transferred to 15°C and incubated overnight before collecting cells by centrifugation and storing pellets at −80°C. Bacterial pellets were thawed, resuspended in 25 ml of lysis buffer (as for RAB-GST proteins), incubated for 30 min in ice, and lysed with a French Press. Lysates were clarified by centrifugation (30,000 x at for 30 min at 4°C and purified in a Ni-Sepharose High Performance column. Imidazole (0.4 M) present in the eluted fraction was removed with a PD-10 column equilibrated in PBS, pH 7.4 containing 5% glycerol and 1 mM DTT. The eluate (3.5 ml) was loaded onto a HiLoad 16/600 Superdex pg column that was run at 1 ml/min. Fractions containing protein were analyzed by SDS-PAGE, stained with Coomassie and pooled as appropriate.

### Total cell extracts

These were carried out as described (Pinar et al., 2019). 70 mg of lyophilized mycelia were ground to a fine powder in 2 ml tubes containing a ceramic bead and a 20 sec pulse of a FastPrep set at power 4. The powder was suspended in 1.5 ml of ‘low KCl buffer’ (25 mM HEPES, pH 7.5, 110 mM KCl, 5 mM MgCl2, 1 mM DTT and 0.1% Triton) containing 10% (v/v) glycerol, complete ULTRA Tablets inhibitor cocktail (Roche) and ∼ 100 μl of 0.6 mm glass beads. The resulting suspension was homogenized with a 15 sec full-power pulse of the FastPrep and proteins were extracted after incubation for 10 min at 4°C in a rotating wheel. This extraction step was repeated two additional times before the resulting homogenate was clarified by centrifugation at 15,000 x g and 4°C in a refrigerated microcentrifuge.

### RAB-GST pull-downs with total cell extracts

6 mg of each extract were mixed with 10 µl of nucleotide-loaded RAB-GST baits in a total volume of 0.4 ml in 0.8 ml Pierce centrifuge columns and the mixtures were incubated for 2 h at 4°C in a rotating wheel. GST-Sepharose beads were collected by low speed centrifugation, washed four times with 0.7 ml of ‘medium KCl buffer’ (25 mM HEPES pH 7.5, 175 mM KCl, 5mM MgCl2, 1 mM DTT and 0,1% Triton X-100) before bound material was eluted with 20 µl of Laemmli loading buffer. 15 µl were run in 7.5% polyacrylamide gels that were analyzed by *α*-HA western blotting and 2µl were run in a 10% polyacrylamide gel for Coomassie staining of the baits.

### RAB-GST pull-downs with purified proteins

Binding reactions were carried out in 0.8 ml Pierce centrifuge columns. Nucleotide-loaded RABs (10 µl of glutathione-Sepharose beads) were mixed with either 2.5 µg of purified UDS1-His6 or with 10 µl of TNT reaction mix primed with appropriate plasmids (pSP64-MyoE, pSP64-HMSV-HA3 or pS64-Uso1-HA3), in 0.4 ml of ‘medium KCl’ buffer containing 10% glycerol. The resulting mix was incubated for 2 h at 4°C in a rotating wheel. Beads were collected by low-speed microcentrifugation and washed four times in the same buffer before eluting bound material with 20 µl of Laemmli loading buffer. 5 µl aliquots were analyzed by western blotting using *α*-His antibody (for UDS1-His6) or *α*-HA antibody (for HMSV-HA3) and 7.5% polyacrylamide gels, or *α*-MyoE antibodies and Biorad’s pre-casted 4-15% polyacrylamide gels (for MyoE).

### Pull-down of the UDS1-HMSV complex with RAB11-GST

10 µl of glutathione-Sepharose beads loaded with RAB11-GST GTP*γ*S or GDP were incubated in Pierce microcolumns for 2 h at 4°C with 2.5 µg of UDS1-His6 and 10 µl of TNT-synthesized HMSV-HA3 in 400 µl of ‘low KCl’ buffer containing 10% glycerol. Beads were washed four times with ‘medium KCl’ buffer. Equal amounts of bound material were analyzed by western blotting using *α*-HA3 and *α*-His antibodies.

### Pull-down with GST-UDS1

GST-UDS1 was purified as described for RAB-GST proteins. 15 µl of glutathione-Sepharose beads containing bait protein fusions were mixed with 10 µl of TNT*®*-synthesized HMSV-HA3 or Uso1-HA3 preys (10 µl of each reaction mix) in 0.4 ml of 25 mM HEPES pH 7.5, 300 mM KCl, 0.5% Triton, 0.5 mM EDTA and 1 mM DTT, using Pierce microcolumns, which were incubated for 2 h at 4°C in a rotating wheel. Beads were washed four times with the same buffer and eluted with 20 µl of Laemmli buffer. 5 µl aliquots were analyzed by *α*-HA western blotting.

### ProtA immunoprecipitations

For *α*-MyoE co-immunoprecipitation experiments of HMSV-HA3, 5 µl samples of Protein A-Sepharose (GE Healthcare) were preincubated with 10 µl each of purified *α*-MyoE or *α*-Uso1 antibodies for 3 h at room temperature. Antibody-loaded beads were mixed with 25 µl of TNT-synthesized MyoE and 25 µl of TNT-synthesized HMSV-HA3 in 0.4 ml of 25 mM HEPES pH 7.5, 500 mM NaCl, 0.5% Triton, 0.5 mM EDTA and 2% BSA, using 0.8 ml Pierce microcolumns. Beads were recovered by microcentrifugation, washed four times in the same buffer (without BSA) and eluted with 20 µl of Laemmli loading buffer. 5 µl of each sample were analyzed by western blot (7.5 % polyacrylamide gel) using *α*- HA mAb. A gel run in parallel was stained with Coomassie blue to asses equal loading of Protein A beads with IgG heavy chains.

### GFP-Trap and western blotting

Cell extracts [strains, MyoE-GFP (MAD4406), UDS1-GFP (MAD6379), HMSV-GFP (MAD7326) and Uso1-GFP (MAD6358)] were prepared as described above, but using the lysis buffer recommended by the manufacturer, which containing 25 mM Tris-HCl pH 7.5, 150 mM NaCl, 0.5 mM EDTA, 0.5% NP40 and Roche’s cOmplete protease inhibitors. Approximately 100 mg of total protein (4 ml of extract) were immunoprecipitated with 25 µl of GFP-Trap magnetic agarose beads (Chromotek gtma-20) following incubation for 2 h at 4°C in a rotating wheel. Beads were washed four times with the same buffer before eluting the immunoprecipitated material with 60 µl of Laemmli buffer. 10 µl aliquots were analyzed by *α*-HA3 western blotting (7.5% polyacrylamide gels) or *α*-MyoE western blotting. 2 µl were analyzed by *α*-GFP western blotting to determine levels of immunoprecipitated baits. Lastly 8 µl were analyzed by SDS-PAGE and silver staining.

### Shotgun proteomic analysis of RAB11-GST effectors

Large scale purification of proteins interacting with the GDP and GTP*γ*S forms of RAB11-GST was carried out as described previously for GST-RAB11 (Pinar & Peñalva, 2017). Bound proteins were loaded onto a 10% polyacrylamide gel, which was run until proteins moved 1 cm into the gel. The protein mixture band was detected by colloidal Coomassie staining, excised and processed for tryptic digestion and subsequent analysis by MS/MS essentially as described (Pinar et al., 2019). For MS/MS analyses of GFP-tagged bait associates, proteins were digested using the ‘on-bead digest protocol for mass spectrometry following immunoprecipitation with Nano-Traps’ recommended by Chromotek. In both cases mass spectra *.raw files were used to search the *A. nidulans* FGSC A4 version_s10m02-r03_orf_trans_allMODI proteome database (8223 protein entries) using Mascot search engine version 2.6 (Matrix Science). Peptides were filtered using Percolator (Kall et al., 2007), with a q-value threshold set to 0.01.

### Analytical ultracentrifugation

Sedimentation equilibrium analysis of UDS1-His was carried out in the Molecular Interactions Facility of the Centro de Investigaciones Biológicas using an XL-A analytical ultracentrifuge (Beckman-Coulter Inc.) equipped with a UV-VIS detector set at 237 nm. Centrifugation was carried out in short (95 µl) columns at speeds ranging from 6000 to 9000 rpm, with a last high-speed (48,000 rpm) run to deplete the protein from the meniscus and obtain the corresponding baseline offsets. Weight-average buoyant molecular weights were determined by fitting, using HeteroAnalysis software (Cole, 2004), a single-species model to the experimental data (corrected for temperature and solvent composition with SEDNTERP software (Laue, 1992)).

### Negative staining electron microscopy

Purified UDS1 was diluted to 0.2 µM in 150 mM NaCl, 25 mM HEPES pH 7.5 and 5% glycerol, and stained with 2% (w/v) uranyl acetate. Specimens were examined under a JEOL 1230 electron microscope equipped with a TVIPS CMOS 4kx4k camera and operated at 100 kV. Data were collected at a nominal magnification of 40,000x, which corresponds to 2.84 Å/pixel at the micrograph level. The length of 71 representative particles selected from multiple micrographs was measured using ImageJ.

### Fluorescence Microscopy

Hyphae were cultured in watch minimal medium (WMM) (Peñalva, 2005). Microscopy chambers, hardware, software and image acquisition procedures have been thoroughly documented (Pinar & Peñalva, 2020), with the sole exception that some of the experiments using the Hamamatsu Gemini beam splitter were carried out in a Leica DMi8 inverted microscope instead of a Leica DMi6000. Z-stacks were deconvolved using Huygens Professional software (Hilversum, Holland), version 20.04.0p5 64 bits. Images (usually MIPs unless otherwise indicated) were contrasted with Metamorph (Molecular Dynamics) and annotated using Corel Draw. Movies were assembled with Metamorph and compressed using QuickTime (Apple Inc.). Quantitation of average MyoE-GFP signals in the SPK was made using MIPs of raw images. Datasets were analyzed with Prism 3.02.

### Statistical analysis

It is described in the legend to Figure 7. Analysis was carried out with GraphPad Prism

## Acknowledgements

We thank Juan M. Luque (Molecular Interactions Facility, Centro de Investigaciones Biológicas) for his help with equilibrium sedimentation analyses, Elena Reoyo for skillful technical assistance and Herb Arst for critical reading of the manuscript. Thanks are due to Spain’s Ministerio de Ciencia e Innovación for grants RTI2018-093344-B100 (MAP) and BFU2017-89143-P (EA-P), as well as for grant-associated Predoctoral contracts BES-2016-077440 (IB.-P/MAP) and PRE2018-086026 (to AG/EA-P). Thanks are also due to the Comunidad de Madrid grant for S2017/BMD-3691 (MAP). Grants were co-funded by European Regional Development and European Social Funds. The authors declare that they do not have any competing financial interests.

## Authors Contributions

MP, AA and IB-P carried out biochemical and genetic experiments. VdR conducted MS/MS analyses, EA-P and AG conducted electron microscopy experiments and MAP carried out fluorescence protein localization analyses, supervised the project and, with MP, wrote the manuscript with input from all authors.

## Supplemental Materials

**Fig. S1.**
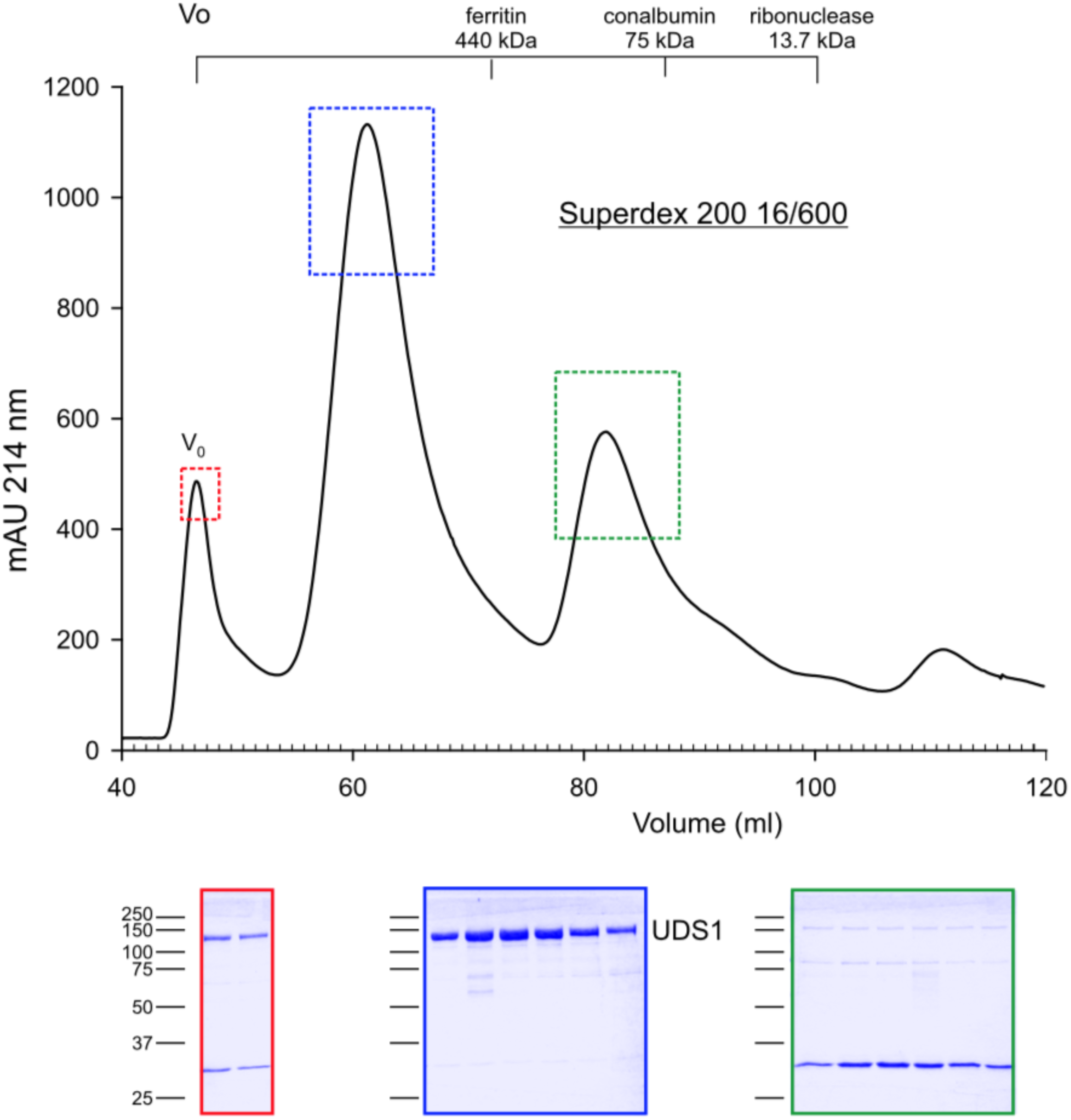
Gel filtration showing the elution profile of UDS1. UDS1-His6 previously purified by Ni2+ chromatography was injected onto a Hiload 16/200 Superdex 600 gel filtration column, which was run at 1ml/min. Aliquots of fractions corresponding to the middle of the peaks were analyzed by SDS-PAGE and Coomassie staining.

**Fig. S2.**
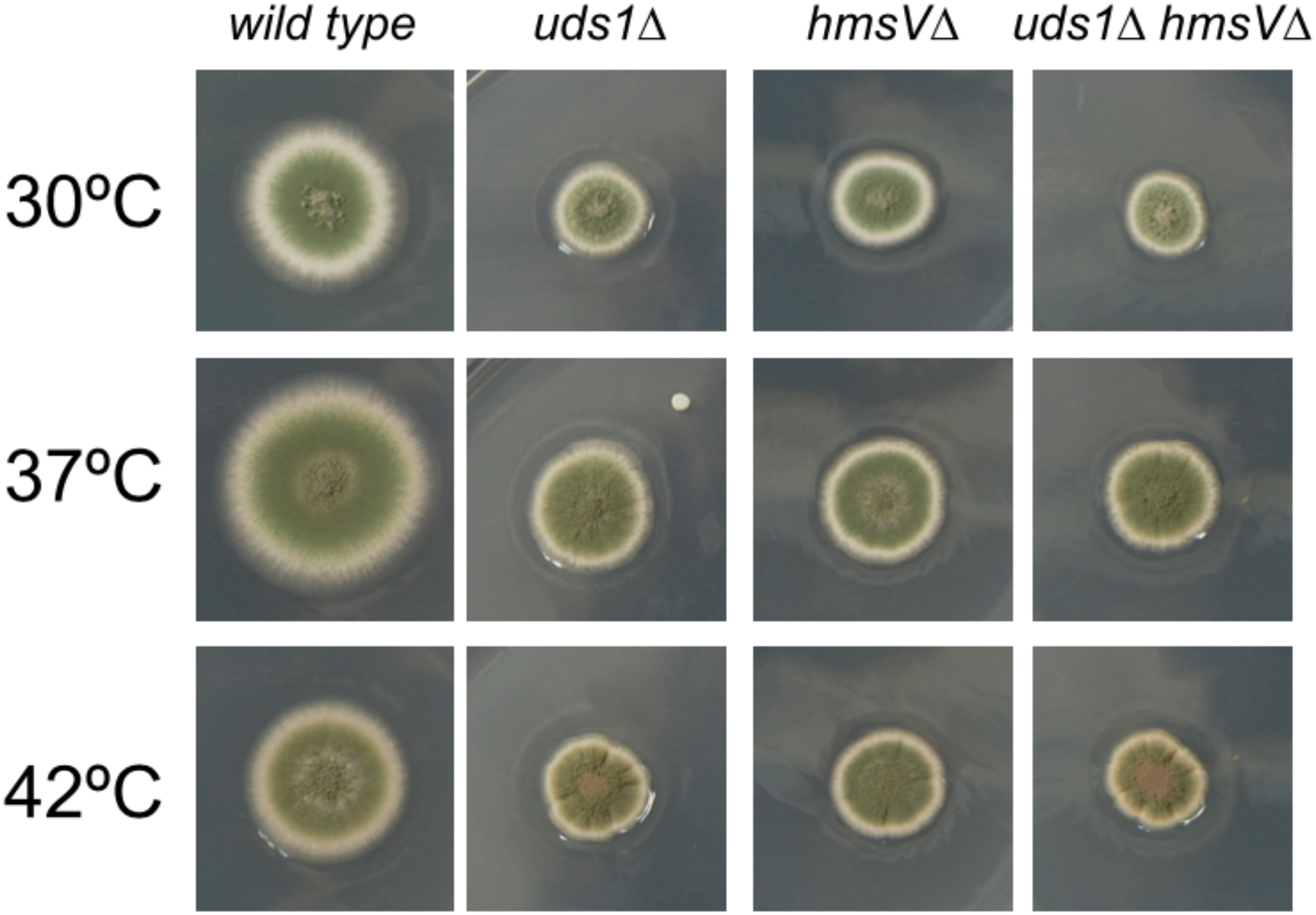
Growth of the double *uds1*Δ *hmsV*Δ mutant is indistinguishable from the single mutants. Growth tests of different strains on *Aspergillus* complete medium at the indicated temperatures.

**Table S1:**
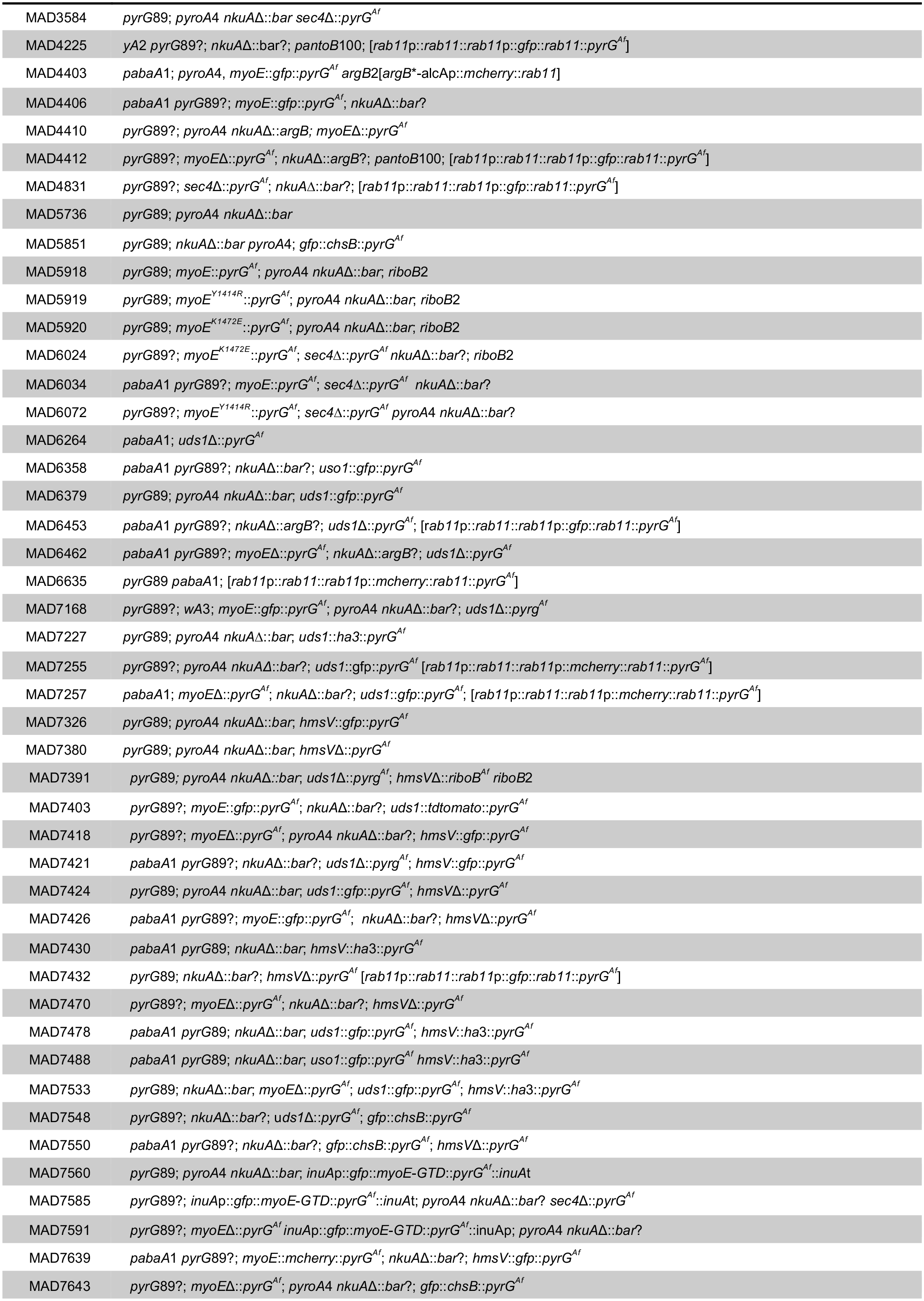
Strain genotypes.

### Legends to Movies

**Movie S1**

GFP-tagged myosin-5 (MyoE) concentrates at the SPK/vesicle supply center of a *sec4*Δ cell. The movie was built with 100 frames representing middle planes of the cell, acquired (streaming to the RAM) at 2.5 fps (total 40 sec). Timestamp is in sec. msec.

**Movie S2**

GFP-GTD (inverted greyscale) arriving at the apical dome of a *myoE*Δ cells by way of MT-dependent transport, implying that the myosin-5 GTD is sufficient to bind SVs. The movie was built with 300 frames representing middle planes of the cell, acquired (streaming to the RAM) at 5 fps (total 1 min). Timestamp is in sec. msec.

**Movie S3**

Co-filming of mCh-RAB11 and UDS1-GFP over a 25 min period. This movie was made with the simultaneously acquired (using a Gemini beam splitter) red and green channel images (shown in inverted greyscale) of a hypha co-expressing both proteins at physiological levels. Built with 101 frames acquired every 15 sec (0.066 fps). Timestamp in min. sec.

**Movie S4**

UDS1-tdT and MyoE-GFP strictly colocalize over time at the SPK. Z-stacks of the corresponding channels (MIPs shown in inverted greyscale) were acquired every 30 sec (0.033 fps) for a total of 38 frames (18.5 min). Timestamp is in min. sec.

**Movie S5**

UDS1-GFP arriving at the apical dome in a *myoE*Δ cell. Note the recurring dots localizing to the plasma membrane, in all likelihood representing SVs containing UDS1 that are being delivered to the dome by MT transport. The movie was made with middle planes streamed to the RAM of the computer every 400 msec (2.5 fps). Timestamp is in sec. msec.

**Movie S6**

mCh-MyoE and HMSV-GFP strictly colocalize over time at the SPK. Z-stacks in the green and red channels were simultaneously acquired with a DualViewer every 15 sec

(0.066 fps) for 15 min. The movie was built with the corresponding MIPs (shown in inverted greyscale). Timestamp is in min.sec.

**Movie S7**

A wild-type and an *hmsV*Δ cell expressing MyoE-GFP are shown alongside using equivalent contrast to underline the reduced levels of MyoE at the mutant’s SPK. Some moving vesicles are visible in the background. The movies were built with 100 middle planes acquired with the same settings and streamed to the computer’s RAM every 100 msec (10 fps) for a total of 10 sec. Timestamp is in sec. msec.

**Movie S8**

One hundred middle planes of a *myoE*Δ cell expressing HMSV-GFP were streamed to the computer’s RAM every 400 msec (2.5 fps) for a total of 40 sec. Note that HMSV distributes like SVs arriving to the dome by MT transport. Time scale is in sec. msec.

## References

Abenza JF, Galindo A, Pantazopoulou A, Gil C, de los Ríos V, Peñalva MA (2010) *Aspergillus* RabB^Rab5^ integrates acquisition of degradative identity with the long-distance movement of early endosomes. Mol Biol Cell 21: 2756–2769

Bielska E, Schuster M, Roger Y, Berepiki A, Soanes DM, Talbot NJ, Steinberg G (2014) Hook is an adapter that coordinates kinesin-3 and dynein cargo attachment on early endosomes. J Cell Biol 204: 989–1007

Cole JL (2004) Analysis of heterogeneous interactions. Methods Enzymol 384: 212–32

Cross JA, Dodding MP (2019) Motor–cargo adaptors at the organelle–cytoskeleton interface. Current Opinion in Cell Biology 59: 16–23

Donovan KW, Bretscher A (2015) Head-to-tail regulation is critical for the in vivo function of myosin V. J Cell Biol 209: 359–365

Goldenring JR (2015) Recycling endosomes. Current Opinion in Cell Biology 35: 117–122

Hales CM, Vaerman J-P, Goldenring JR (2002) Rab11 Family Interacting Protein 2 Associates with Myosin Vb and Regulates Plasma Membrane Recycling. J Biol Chem 277: 50415–50421

Hammer JA, 3rd, Sellers JR (2012) Walking to work: roles for class V myosins as cargo transporters. Nat Rev Mol Cell Biol 13: 13–26

Hernández-González M, Bravo-Plaza I, Pinar M, de los Ríos V, Arst HN, Jr., Peñalva MA (2018a) Endocytic recycling via the TGN underlies the polarized hyphal mode of life. PLoS Genetics 14: e1007291

Hernández-González M, Pantazopoulou A, Spanoudakis D, Seegers CLC, Peñalva MA (2018b) Genetic dissection of the secretory route followed by a fungal extracellular glycosyl hydrolase. Mol Microbiol 109: 781–800

Hodges AR, Bookwalter CS, Krementsova EB, Trybus KM (2009) A nonprocessive class V myosin drives cargo processively when a kinesin-related protein is a passenger. Curr Biol 19: 2121–5

Hohmann-Marriott MF, Uchida M, van de Meene AM, Garret M, Hjelm BE, Kokoori S, Roberson RW (2006) Application of electron tomography to fungal ultrastructure studies. New Phytol 172: 208–220

Ikebe C, Konishi M, Hirata D, Matsusaka T, Toda T (2011) Systematic localization study on novel proteins encoded by meiotically up-regulated ORFs in fission yeast. Biosci Biotechnol Biochem 75: 2364–70

Jin Y, Sultana A, Gandhi P, Franklin E, Hamamoto S, Khan AR, Munson M, Schekman R, Weisman LS (2011) Myosin V transports secretory vesicles via a Rab GTPase cascade and interaction with the exocyst complex. Dev Cell 21: 1156–70

Kall L, Canterbury JD, Weston J, Noble WS, MacCoss MJ (2007) Semi-supervised learning for peptide identification from shotgun proteomics datasets. Nat Methods 4: 923–5

Laue TM, Shah, B.D., Ridgeway, T.M. and Pelletier, S.L (1992) Interpretation of analytical sedimentation data for proteins. In Analytical Ultracentrifugation in Biochemistry and Polymer Science Harding SE, Rowe, A.J. and Horton, J.C. (ed) pp 90–125. Cambridge: Royal Society of Chemistry

Li BX, Satoh AK, Ready DF (2007) Myosin V, Rab11, and dRip11 direct apical secretion and cellular morphogenesis in developing Drosophila photoreceptors. J Cell Biol 177: 659–669

Lipatova Z, Tokarev AA, Jin Y, Mulholland J, Weisman LS, Segev N (2008) Direct interaction between a myosin V motor and the Rab GTPases Ypt31/32 is required for polarized secretion. Mol Biol Cell 19: 4177–87

Lwin KM, Li D, Bretscher A (2016) Kinesin-related Smy1 enhances the Rab-dependent association of myosin-V with secretory cargo. Mol Biol Cell

Matsui T, Ohbayashi N, Fukuda M (2012) The Rab interacting lysosomal protein (RILP) homology domain functions as a novel effector domain for small GTPase Rab36: Rab36 regulates retrograde melanosome transport in melanocytes. J Biol Chem 287: 28619–31

Nayak T, Szewczyk E, Oakley CE, Osmani A, Ukil L, Murray SL, Hynes MJ, Osmani SA, Oakley BR (2005) A versatile and efficient gene targeting system for *Aspergillus nidulans*. Genetics 172: 1557–1566

Pantazopoulou A, Pinar M, Xiang X, Peñalva MA (2014) Maturation of late Golgi cisternae into RabE^RAB11^ exocytic post-Golgi carriers visualized *in vivo*. Mol Biol Cell 25: 2428–2443

Pashkova N, Jin Y, Ramaswamy S, Weisman LS (2006) Structural basis for myosin V discrimination between distinct cargoes. EMBO J 25: 693–700

Peñalva MA (2005) Tracing the endocytic pathway of *Aspergillus nidulans* with FM4-64. Fungal Genet Biol 42: 963–975

Peñalva MA, Zhang J, Xiang X, Pantazopoulou A (2017) Transport of fungal RAB11 secretory vesicles involves myosin-5, dynein/dynactin/p25 and kinesin-1 and is independent of kinesin-3. Mol Biol Cell 28: 947–961

Pfeffer SR (2013) Rab GTPase regulation of membrane identity. Curr Opin Cell Biol 25: 414–9

Pinar M, Arias-Palomo E, de los Ríos V, Arst HN, Jr., Peñalva MA (2019) Characterization of *Aspergillus nidulans* TRAPPs uncovers unprecedented similarities between fungi and metazoans and reveals the modular assembly of TRAPPII. PLOS Genetics 15: e1008557

Pinar M, Arst HN, Jr., Pantazopoulou A, Tagua VG, de los Ríos V, Rodríguez-Salarichs J, Díaz JF, Peñalva MA (2015) TRAPPII regulates exocytic Golgi exit by mediating nucleotide exchange on the Ypt31 orthologue RabE/RAB11. Proc Natl Acad Sci USA 112: 4346–4351

Pinar M, Peñalva MA (2017) *Aspergillus nidulan*s BapH is a RAB11 effector that connects membranes in the Spitzenkörper with basal autophagy. Mol Microbiol 106: 452–468

Pinar M, Peñalva MA (2020) *En bloc* TGN recruitment of *Aspergillus* TRAPPII reveals TRAPP maturation as unlikely to drive RAB1-to-RAB11 transition. J Cell Sci 133: jcs241141

Pinar M, Peñalva MA (2021) The fungal RABOME: RAB GTPases acting in the endocytic and exocytic pathways of *Aspergillus nidulans* (with excursions to other filamentous fungi). *Mol Microbiol in press*

Pylypenko O, Attanda W, Gauquelin C, Lahmani M, Coulibaly D, Baron B, Hoos S, Titus MA, England P, Houdusse AM (2013) Structural basis of myosin V Rab GTPase-dependent cargo recognition. Proceedings of the National Academy of Sciences 110: 20443–20448

Pylypenko O, Welz T, Tittel J, Kollmar M, Chardon F, Malherbe G, Weiss S, Michel CIL, Samol-Wolf A, Grasskamp AT, Hume A, Goud B, Baron B, England P, Titus MA, Schwille P, Weidemann T, Houdusse A, Kerkhoff E (2016) Coordinated recruitment of Spir actin nucleators and myosin V motors to Rab11 vesicle membranes. eLife 5: e17523

Qiu R, Zhang J, Xiang X (2019) LIS1 regulates cargo-adapter–mediated activation of dynein by overcoming its autoinhibition in vivo. J Cell Biol 218: 3630–3646

Randall TS, Yip YY, Wallock-Richards DJ, Pfisterer K, Sanger A, Ficek W, Steiner RA, Beavil AJ, Parsons M, Dodding MP (2017) A small-molecule activator of kinesin-1 drives remodeling of the microtubule network. Proc Natl Acad Sci USA 114: 13738–13743

Roland JT, Bryant DM, Datta A, Itzen A, Mostov KE, Goldenring JR (2011) Rab GTPase–Myo5B complexes control membrane recycling and epithelial polarization. Proc Natl Acad Sci USA 108: 2789–2794

Santiago-Tirado FH, Legesse-Miller A, Schott D, Bretscher A (2011) PI4P and Rab inputs collaborate in myosin-V-dependent transport of secretory compartments in yeast. Dev Cell 20: 47–59

Schafer JC, Baetz NW, Lapierre LA, McRae RE, Roland JT, Goldenring JR (2014) Rab11-FIP2 interaction with MYO5B regulates movement of Rab11a-containing recycling vesicles. Traffic 15: 292–308

Schuchardt I, Aßmann D, Thines E, Schuberth C, Steinberg G (2005) Myosin-V, Kinesin-1, and Kinesin-3 Cooperate in Hyphal Growth of the Fungus *Ustilago maydis*. Mol Biol Cell 16: 5191–5201

Sckolnick M, Krementsova EB, Warshaw DM, Trybus KM (2013) More than just a cargo adapter, melanophilin prolongs and slows processive runs of myosin Va. J Biol Chem 288: 29313–22

Sharpless KE, Harris SD (2002) Functional characterization and localization of the *Aspergillus nidulans* formin SEPA. Mol Biol Cell 13: 469–479

Szewczyk E, Nayak T, Oakley CE, Edgerton H, Xiong Y, Taheri-Talesh N, Osmani SA, Oakley BR (2006) Fusion PCR and gene targeting in *Aspergillus nidulans*. Nat Protoc 1: 3111–20

Taheri-Talesh N, Xiong Y, Oakley BR (2012) The Functions of Myosin II and Myosin V Homologs in Tip Growth and Septation in *Aspergillus nidulans*. PLoS ONE 7: e31218

Tilburn J, Scazzocchio C, Taylor GG, Zabicky-Zissman JH, Lockington RA, Davies RW (1983) Transformation by integration in *Aspergillus nidulans*. Gene 26: 205–211

Todd RB, Davis MA, Hynes MJ (2007) Genetic manipulation of Aspergillus nidulans : meiotic progeny for genetic analysis and strain construction. Nat Protoc 2: 811–821

Wang Z, Edwards JG, Riley N, Provance DW, Karcher R, Li X-d, Davison IG, Ikebe M, Mercer JA, Kauer JA, Ehlers MD (2008) Myosin Vb Mobilizes Recycling Endosomes and AMPA Receptors for Postsynaptic Plasticity. Cell 135: 535–548

Wei Z, Liu X, Yu C, Zhang M (2013) Structural basis of cargo recognitions for class V myosins. Proc Natl Acad Sci USA 110: 11314–9

Wong S, Hepowit NL, Port SA, Yau RG, Peng Y, Azad N, Habib A, Harpaz N, Schuldiner M, Hughson FM, MacGurn JA, Weisman LS (2020) Cargo Release from Myosin V Requires the Convergence of Parallel Pathways that Phosphorylate and Ubiquitylate the Cargo Adaptor. Curr Biol 30: 4399–4412.e7

Wong S, Weisman LS (2021) Roles and regulation of myosin V interaction with cargo. Adv Biol Regul: 100787

Wu S-Z, Bezanilla M (2018) Actin and microtubule cross talk mediates persistent polarized growth. J Cell Biol 217: 3531–3544

Wu XS, Rao K, Zhang H, Wang F, Sellers JR, Matesic LE, Copeland NG, Jenkins NA, Hammer JA, 3rd (2002) Identification of an organelle receptor for myosin-Va. Nat Cell Biol 4: 271–8

Wu XS, Tsan GL, Hammer JA, III (2005) Melanophilin and myosin Va track the microtubule plus end on EB1. J Cell Biol 171: 201–207

Yao X, Wang X, Xiang X (2014) FHIP and FTS proteins are critical for dynein-mediated transport of early endosomes in Aspergillus. Mol Biol Cell 25: 2181–2189

Zhang J, Qiu R, Arst HN, Jr., Peñalva MA, Xiang X (2014) HookA is a novel dynein-early endosome linker critical for cargo movement in vivo. J Cell Biol 204: 1009–26

Zhang J, Tan K, Wu X, Chen G, Sun J, Reck-Peterson SL, Hammer JA, 3rd, Xiang X (2011) *Aspergillus* Myosin-v supports polarized growth in the absence of microtubule-based transport. PLoS ONE 6: e28575

Zheng P, Nguyen TA, Wong JY, Lee M, Nguyen TA, Fan JS, Yang D, Jedd G (2020) Spitzenkorper assembly mechanisms reveal conserved features of fungal and metazoan polarity scaffolds. Nat Commun 11: 2830

